# Defective RNA Polymerase III is negatively regulated by the SUMO-Ubiquitin-Cdc48 Pathway

**DOI:** 10.1101/259846

**Authors:** Zheng Wang, Catherine Wu, Aaron Aslanian, John R. Yates, Tony Hunter

## Abstract

Transcription by RNA polymerase III (Pol III) is an essential cellular process, and mutations in Pol III can cause neurodegenerative disease in humans. However, in contrast to Pol II transcription, which has been extensively studied, the knowledge of how Pol III is regulated is very limited. We report here that in budding yeast, *Saccharomyces cerevisiae*, Pol III is negatively regulated by the <u>S</u>mall <u>U</u>biquitin-like <u>MO</u>difier (SUMO), an essential post-translational modification pathway. Besides sumoylation, Pol III is also targeted by ubiquitylation and the Cdc48/p97 segregase, the three of which likely act in a sequential manner and eventually lead to proteasomal degradation of Pol III subunits, thereby repressing Pol III transcription. This study not only uncovered a regulatory mechanism for Pol III, but also suggests that the SUMO and ubiquitin modification pathways and the Cdc48/p97 segregase can be potential therapeutic targets for Pol III-related human diseases.

## INTRODUCTION

Eukaryotes have three conserved DNA-directed RNA polymerases (RNA Pols) (Roeder & Rutter, 1969, 1970; Weinmann & Roeder, 1974; Zylber & Penman, 1971), where Pol I transcribes most of the rRNAs, Pol II transcribes mRNA, and Pol III transcribes tRNA, 5S rRNA, as well as some non-coding RNAs, such as the U6 snRNA involved in mRNA splicing. The Pol III machinery includes the polymerase itself (composed of 17 subunits), as well as basal transcription factors TFIIIA, the TFIIIB complex, and the TFIIIC complex (Geiduschek & Kassavetis, 2001). In budding yeast, *Saccharomyces cerevisiae*, TFIIIB is composed of Brf1, Bdp1, and TBP. TFIIIC is composed of Tfc1, Tfc3, Tfc4, Tfc6, Tfc7, and Tfc8. For 5S rRNA transcription, all three basal transcription factors are required, whereas tRNA transcription only requires TFIIIB and TFIIIC. As important as it is for normal cell physiology, Pol III plays critical roles in pathological processes, such as virus infection (Chiu, Macmillan, & Chen, 2009) and tumorigenesis (White, 2004). In addition, Pol III mutations were recently found to cause neurodegenerative diseases in humans. Mutations that cause hypomyelinating leukodystrophy with 4 H syndrome occur predominantly in the largest two subunits of Pol III, POLR3A and POLR3B (Rpc160 and Rpc128 in yeast, respectively) (Bernard et al., 2011; Saitsu et al., 2011; Shimojima et al., 2014; Synofzik, Bernard, Lindig, & Gburek-Augustat, 2013; Terao et al., 2012; Tetreault et al., 2011), with a few in POLR1C (Rpc40 in yeast) (Thiffault et al., 2015), a subunit shared by Pol I and Pol III. Four mutations in BRF1 were found to cause a cerebellar-facial-dental syndrome (Borck et al., 2015).

How Pol III transcription is regulated is still poorly understood. Current knowledge of Pol III regulation is largely limited to phosphorylation of the Pol III machinery components, such as Maf1 (Moir et al., 2006; Oficjalska-Pham et al., 2006; Roberts, Wilson, Huff, Stewart, & Cairns, 2006) and the Rpc53 subunit of Pol III (Lee, Moir, McIntosh, & Willis, 2012). Maf1 is a robust Pol III repressor (Boguta, 2013; Moir & Willis, 2013). Upon stress, Maf1 is dephosphorylated and translocated into the nucleus, where it binds Pol III and blocks its interaction with TFIIIB (Desai et al., 2005; Moir et al., 2006; Roberts et al., 2006; Vannini et al., 2010). Phosphorylation of Rpc53 by the Mck1 and Kns1 kinases also represses Pol III under stress conditions, although the mechanism is unclear (Lee et al., 2012). SUMO is another potential regulator for Pol III that could act as a transcriptional repressor (Neyret-Kahn et al., 2013; Rohira, Chen, Allen, & Johnson, 2013), or activator (Chymkowitch et al., 2017), but how sumoylation regulates Pol III is largely unclear. Therefore, deeper insights regarding the regulatory mechanisms of Pol III transcription are needed to design therapeutic tools that can be used to modulate Pol III activity accordingly in human diseases.

Post-translational modification by SUMO is a conserved pathway and is essential for viability in most organisms (Kerscher, Felberbaum, & Hochstrasser, 2006). Similar to ubiquitin modification, SUMO is conjugated to a lysine residue within the target protein through a cascade of reactions catalyzed by a SUMO-specific E1 activating enzyme (Johnson, Schwienhorst, Dohmen, & Blobel, 1997), an E2 conjugating enzyme (Johnson & Blobel, 1997), and E3 ligases (Johnson & Gupta, 2001; Strunnikov, Aravind, & Koonin, 2001; Takahashi, Kahyo, Toh, Yasuda, & Kikuchi, 2001; Zhao, Wu, & Blobel, 2004). SUMO proteases are responsible for both the maturation of the SUMO polypeptide (Li & Hochstrasser, 1999) and the removal of SUMO from modified proteins (Li & Hochstrasser, 2000). Sumoylation can trigger ubiquitylation through the activity of <u>S</u>UMO-<u>T</u>argeted <u>Ub</u>iquitin E3 <u>L</u>igases (STUbLs) (Mullen & Brill, 2008; Prudden et al., 2007; Sun, Leverson, & Hunter, 2007; Xie et al., 2007). Besides ubiquitylation, SUMO can also recruit the Cdc48 (p97)-Ufd1-Npl4 segregase complex, through the SUMO-interacting motifs (SIMs) in Cdc48 and Ufd1 (Bergink et al., 2013; Nie et al., 2012). The key to fully understand the functions of sumoylation is its substrates. Ever since its discovery two decades ago (Mahajan, Delphin, Guan, Gerace, & Melchior, 1997; Matunis, Coutavas, & Blobel, 1996; Okura et al., 1996), biochemical approaches have been greatly improved to identify thousands of sumoylated proteins, as well as their conjugation sites (Hendriks et al., 2014; Lamoliatte et al., 2014; Tammsalu et al., 2014), underscoring the importance of this modification in the cell. However, how sumoylation affects the functions of its protein substrates is still a challenging question that remains largely unanswered, because mutating the conjugation sites usually does not cause any obvious phenotype. To address this issue, a phenotype-based genetic method is needed.

## RESULTS

### A reverse suppressor screen identified Pol III as a major functional target of SUMO

We designed a reverse suppressor screen in budding yeast, *Saccharomyces cerevisiae*, with the goal of identifying proteins or pathways, for which loss of sumoylation results in a phenotype. Specifically, the screen looks for lethal or sick mutations that can be rescued by a dominant Q56K mutation in SUMO (*SMT3-Q56K*). *SMT3-Q56K* is one of the SUMO pathway mutations identified previously in the *mot1-301* suppressor screen (Z. Wang, Jones, & Prelich, 2006) (Table S1), and it suppresses *mot1-301* dominantly (Figure S1). *SMT3-Q56K* cells are viable, suggesting the mutated protein is partially functional (data not shown). To perform the screen (Figure S2, *Materials and Methods*), yeast cells were first transformed with a plasmid carrying *URA3* and *SMT3-Q56K*, followed by random mutagenesis, and allowed to grow into single colonies. Using the *ade2/ade3* color assay, yeast cells will turn red in the presence of the plasmid. If a clone carries a lethal/sick mutation that can be rescued by *SMT3-Q56K*, the cells can no longer lose the plasmid in order to grow. All cells from this clone will thus maintain the plasmid, forming a colony that is uniformly red. Such colonies will be sensitive to 5-fluoroorotic acid (5FOA), which counter-selects the *URA3* gene on the plasmid. Mutated genes can subsequently be cloned by transforming with a genomic DNA library and screening for 5FOA-resistant colonies. Mutations are then identified by PCR sequencing of the gene locus.

The screen results are summarized in Table 1. First, the screen identified mutations in expected genes, including *MOT1* and *SMT3.* The screen also revealed mutations in *AOS1* (SUMO E1) and *ULP2* (SUMO protease), which was not surprising, as they encode enzymes in the SUMO pathway. Strikingly, the remaining 13 mutations were all in genes encoding components of the Pol III transcription machinery, including the largest two subunits of Pol III (Rpc160 and Rpc128), a TFIIIB subunit (Brf1), and two TFIIIC subunits (Tfc1 and Tfc6). To confirm the screen results, the identified mutations were introduced into wild type cells, and subsequently crossed with a strain lacking a major SUMO E3 ligase, Siz1 *(siz1A).* For example, introduction of the M809I mutation in Rpc160 (Figure 1A) and the A704T mutation in Rpc128 (Figure 1B) caused severe growth defects, while *siz1Δ* fully rescued *rpc160-M809I* and partially rescued *rpc128-A704T*, as expected. However, the deletion of the closely related SUMO E3 ligase, *SIZ2*, did not rescue (Figure 1C). Correlating with the growth phenotype, the amount of total tRNA (Figure 1D) as well as individual tRNA species, including mature and pre-mature intron-containing tRNAs (Figure 1E), were dramatically decreased in *rpc160* mutant cells, but were restored to normal levels by *siz1Δ.* Interestingly, *siz1Δ* did not further increase tRNA levels in wild type *RPC160* cells. No change in 5S rRNA was observed, which is a common phenomenon. This is likely because 5S rRNA is produced in excess, and that Pol III has a higher affinity for the initiation complex containing TFIIIA, which is required for the transcription of 5S rRNA, but not tRNAs (Stettler, Mariotte, Riva, Sentenac, & Thuriaux, 1992).

**Table 1.** Summary of mutations rescued by *SMT3-Q56K.*

| Gene | Protein | # of alleles | Mutations |
| --- | --- | --- | --- |
| *MOT1* | Negative regulator of TBP | 3 | *mot1-399* (Q1587 Stop)<br>*mot1-517* (G1410R)<br>*mot1-753* (G1300S) |
| *SMT3* | SUMO | 4 | Not sequenced |
| *AOS1* | SUMO E1 | 1 | *aos1-492* (G56S) |
| *ULP2* | SUMO protease | 4 | *ulp2-4* (S108 Stop)<br>*ulp2-253* (G265D)<br>*ulp2-527* (W532 Stop)<br>*ulp2-63* (W532 Stop) |
| *RPC160* | RNA Pol III subunit | 8 | *rpc160-58* (M809I)<br>*rpc160-85* (G1297D)<br>*rpc160-33* (T379I)<br>*rpc160-419* (A880T)<br>*rpc160-426* (E282K)<br>*rpc160-480* (G1098D)<br>*rpc160-628* (R365K)<br>*rpc160-211* (G606S) |
| *RPC128* | RNA Pol III subunit | 2 | *rpc128-202* (A704T)<br>*rpc128-578* (D501N) |
| *BRF1* | TFIIIB subunit | 1 | *brf1-137* (S271L) |
| *TFC1* | TFIIIC subunit | 1 | *tfc1-321* (N255K, Fs)<br>(AAC-AAAC, Ins, Fs) |
| *TFC6* | TFIIIC subunit | 1 | *tfc6-192* (G391E) |

**Figure 1.**
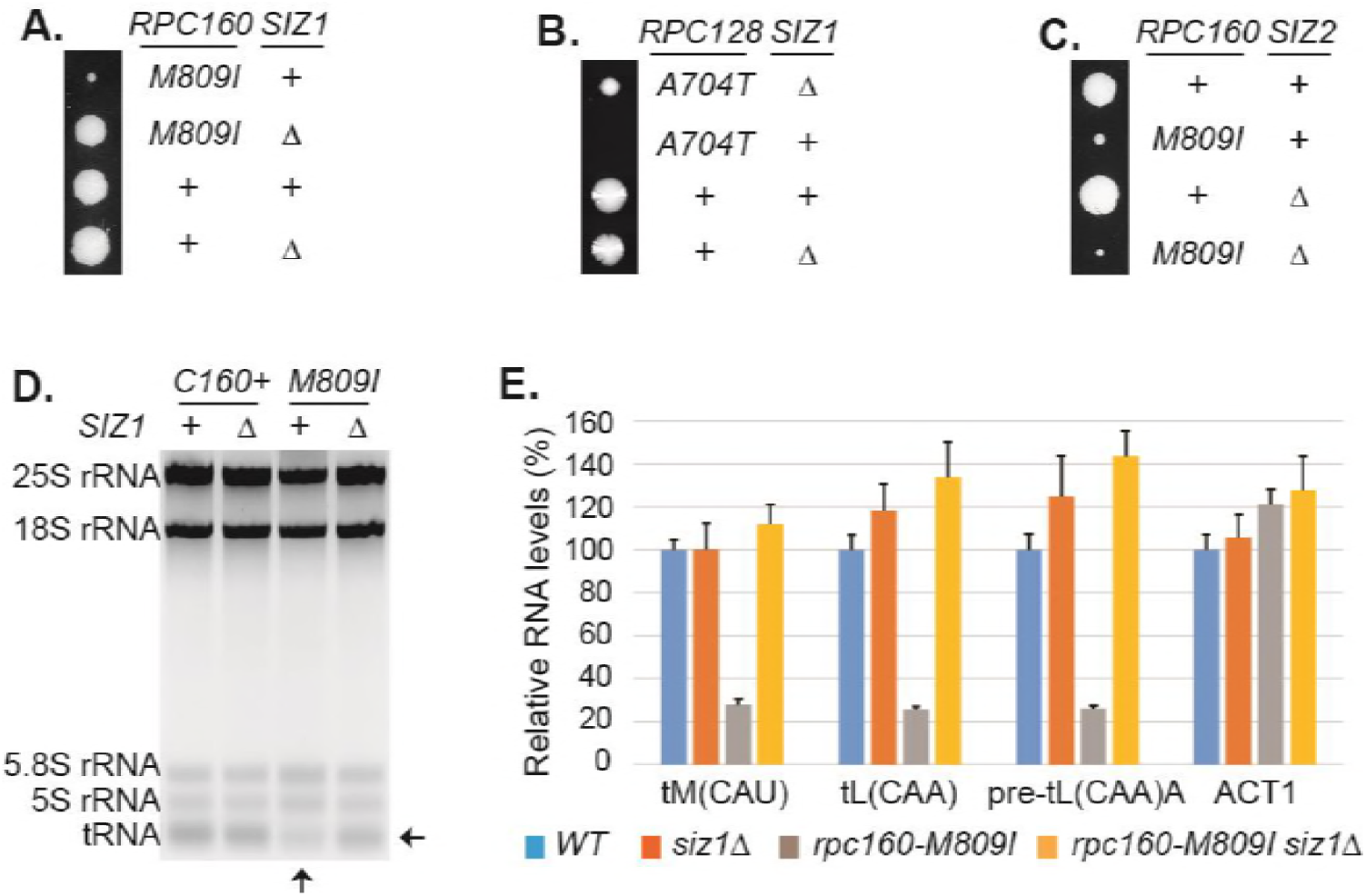
Disrupting sumoylation rescues Pol III mutations. **(A)** Tetrad analysis of a cross between *rpc160-M809I* and *siz1A* Tetrads were dissected on YPD, then incubated at 30°C for four days. The offspring of one representative tetrad was shown with genotypes labeled. **(B)** Similar tetrad analysis for *rpc128-A704T* and *siz1Δ* **(C)** Similar tetrad analysis for *rpc160-M809I* and *siz2Δ* **(D)** 2 µg of RNA extracted from the indicated strains was run on a 2.8% agarose gel containing ethidium bromide, then visualized with UV. **(E)** RNA from (C) was reverse transcribed into cDNA, followed by real time PCR analysis. GAPDH transcripts were used as loading control. Data are mean ± standard deviation calculated from 6 data points (two biological replicates and three technical replicates), presented as relative amount compared to wild type. The intron-containing pre-mature tRNA (pre-tL(CAA)A) is short-lived, so its abundance reflects the Pol III transcriptional activity.

Based on the Cryo-EM structure of Pol III (Hoffmann et al., 2015), most of the mutations identified by the screen occurred on residues close to the bound DNA template or the growing RNA chain (Table S2), suggesting that the mutations could impact Pol III enzyme activity and thus cause severe growth defects. However, none of these mutants have been reported before, so their actual enzymatic defects are unclear. We therefore tested whether previously described Pol III mutations, including *rpc31-236* which is an initiation defective mutant (Thuillier, Stettler, Sentenac, Thuriaux, & Werner, 1995), as well as two elongation mutants, *rpc 160–112* (Dieci et al., 1995) and *rpc160-270* (Thuillier, Brun, Sentenac, & Werner, 1996), could be rescued by reduced sumoylation, and found that all three mutants were rescued by *siz1Δ* (Figure S3A). Besides these loss-of-function mutations, when expression of wild type *RPC 160* was reduced by growth of a strain in which the only *RPC160* gene was under the *GAL1* promoter in glucose, the resultant slow growth phenotype was partially rescued by *siz1Δ* (Figure S3B). We also tested whether human Pol III mutations that cause neuronal diseases, which were introduced into yeast Rpc160, Rpc128, and Brf1 at corresponding positions based on sequence homology, could be rescued when sumoylation was compromised. Among the seventeen Rpc160 mutations tested for growth under normal conditions and at elevated temperature (37°C) (Table S3), two single (Q608K and E1329K) and two double mutations (D384N, N789I and Q608K, G1308S) caused growth defects, which could all be rescued by *siz1Δ*, except for E1329K (Figure S3C). For Rpc 128, only one of the five single mutations (L1027P) showed slower growth, which was rescued by *siz1Δ* (Figure S3D). All *rpc128* double mutations were lethal, and not rescued by *siz1A* (Table S3). For the four *brf1* single mutations, three showed growth defects, two of which were rescued by *siz1Δ* (Figure S3E). These results confirmed the roles of SUMO in Pol III transcription, suggesting that SUMO can repress Pol III but the effect is most obvious when Pol III activity is greatly reduced either through decreased expression or inactivating mutation.

### SUMO preferentially targets Pol III and acts independently of the known Pol III repressors

To gain further insights about the functions of SUMO in Pol III transcription, several specificity tests were performed. It is surprising that our screen only identified Pol III but not either of the other two polymerases, given the fact that the three polymerases are very similar to each other, with many related subunits and even shared subunits. We therefore first tested if it was due to a specific function of SUMO or simply because the screen was not saturated, by introducing similar mutations into the three RNA polymerases, such as an aspartic acid mutation to the glycine residue in the highly conserved “trigger loop” domain in the largest subunits of the polymerases (Fernandez-Tornero et al., 2013; Hoffmann et al., 2015; D. Wang, Bushnell, Westover, Kaplan, & Kornberg, 2006) (Figure 2A). Interestingly, while the G to D mutation caused severe growth defect in all three cases, only the G1098D mutation in Rpc160 (Pol III) was rescued by *siz1Δ*, suggesting SUMO preferentially targets Pol III rather than Pol I or Pol II.

**Figure 2.**
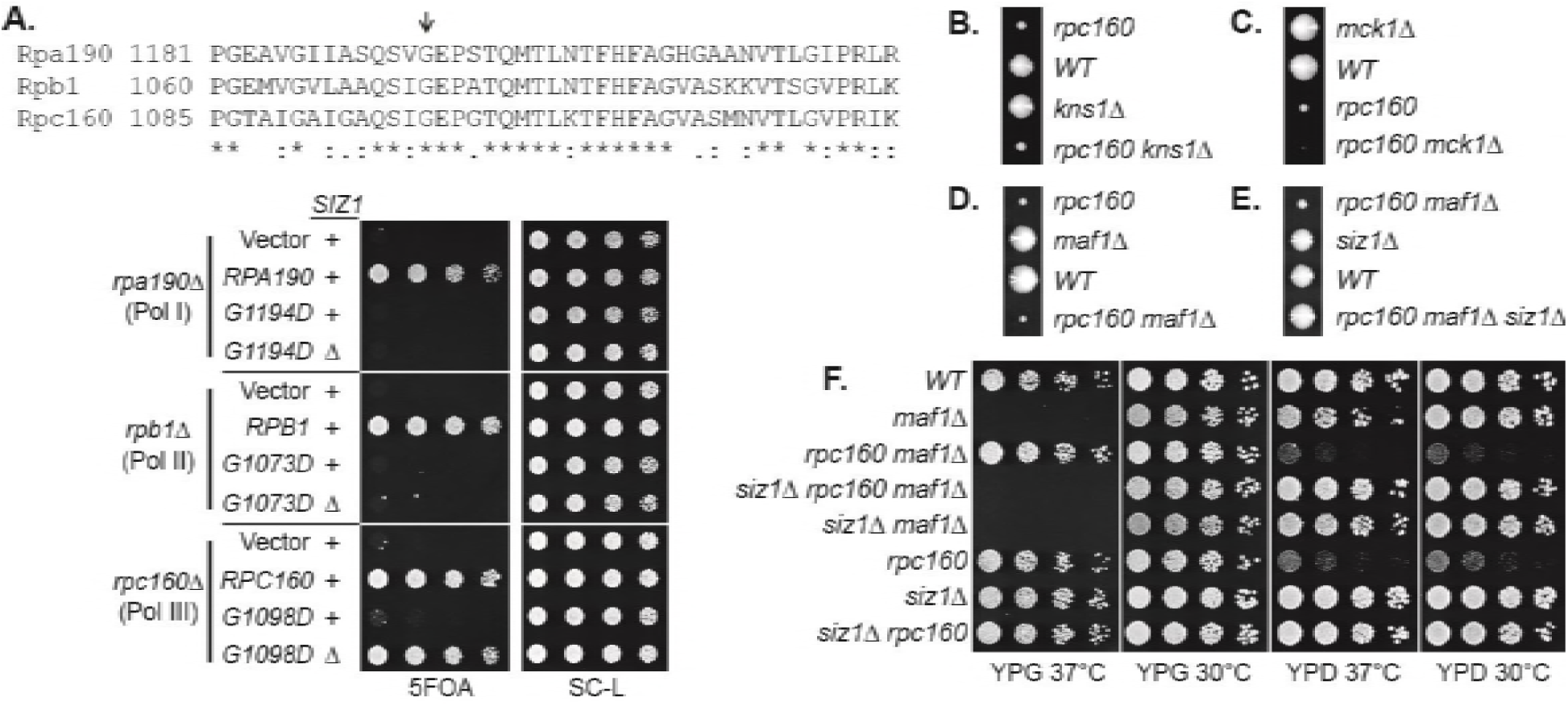
Specificity of the rescue effect. **(A)** The *rpa190A, rpb1A*, or *rpc160Δ* strain carries a *URA3* plasmid carrying wild type *RPA190, RPB1*, and *RPC 160* gene, respectively, in order to maintain viability. These strains were then transformed with *LEU2* plasmids carrying the indicated wild type or mutant alleles, and selected on synthetic media lacking leucine (SC-L). Transformants were spotted in 5-fold serial dilutions onto a 5FOA plate to assess the growth phenotype of the mutant allele, as the original *URA3* plasmids were shuffled out of the cell in the presence of 5FOA. **(B-D)** Tetrad analysis between *rpc160-M809I* (shown as *rpc160)* and *kns1* ***A****, mck1* ***A***, and *maf1* ***A*. (E)** Tetrad analysis between *rpc 160-M809I siz1Δ* and *maf1* ***A*. (F)** The indicated strains from the cross in (E) were plated in 5-fold dilutions onto YPD (glucose) or YPG (glycerol) plates and incubated at 30°C or 37°C as indicated.

We next compared SUMO to known Pol III repressors, including Maf1 and the Mck1 and Kns1 kinases. Surprisingly, none of these proteins, when depleted by deleting the encoding genes, could rescue the *rpc160* mutant growth defect (Figure 2B, 2C, **and** 2D). Furthermore, *siz1Δ* could rescue *rpc160* even in the absence of Maf1 (Figure 2E), and reverse the ability of *rpc 160* to rescue *maf1 Δ* on glycerol media (Figure 2F). Therefore, SUMO specifically targets Pol III for repression, and it does so through a mechanism that is independent of Maf1 or the Mck1 and Kns1 kinases.

### SUMO represses Pol III by modifying Rpc53

To understand the underlying molecular mechanism, the key is the relevant sumoylated protein(s), which is likely in the Pol III machinery itself. To identify this protein(s), we made a strain expressing Flag-tagged Rpc160-M809I and GFP-SUMO, as well as single-tagged strains as negative controls. Mutant *rpc160-M809I was* used, because sumoylation has stronger effects on mutant Pol III than the wild type (Figure 1D **and** 1E). Pol III was first immunoprecipitated (IP) by anti-Flag beads (Figure 3A). Clear sumoylation signals were detected associated with Pol III. These Pol III-associated sumoylated proteins were released and subsequently purified by IP with GFP-trap beads, then analyzed by mass-spectrometry (Figure 3B). Four Pol III components (Rpc160, Rpc82, Rpc53, and Rpc37) were identified. Rpc160 (~160 kDa) is likely to be a contaminant, because the detected sumoylated species ran no slower than the 150 kDa marker band (Figure 3A). Rpc82 sumoylation was reported previously to occur on K406 (Panse, Hardeland, Werner, Kuster, & Hurt, 2004). However, *rpc82-K406R* did not rescue *rpc160-M809I* (data not shown), suggesting Rpc82 is not the relevant sumoylated protein.

**Figure 3.**
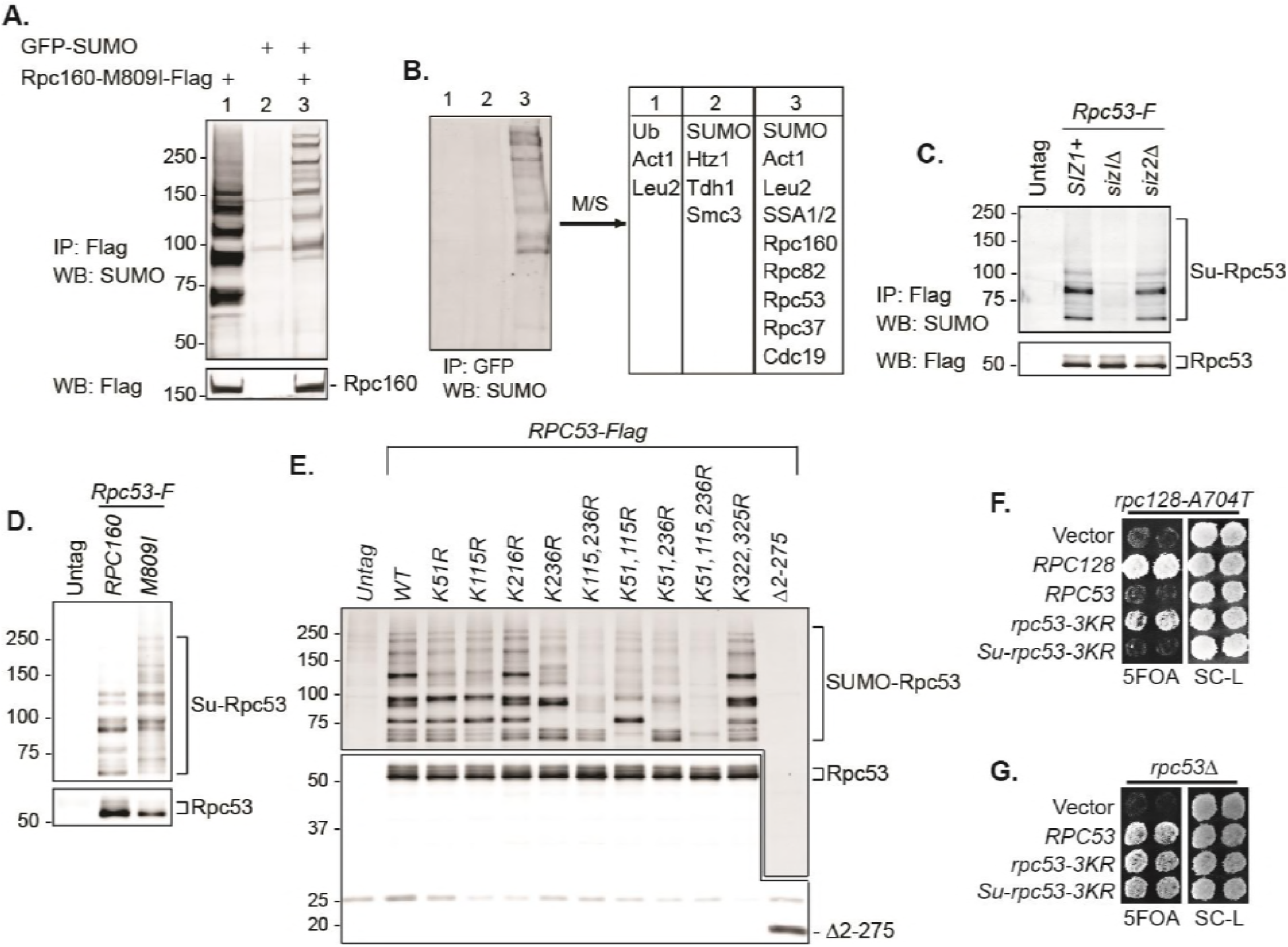
SUMO represses Pol III by modifying Rpc53. **(A)** Total protein extracted from the indicated strains was subjected to anti-Flag IP to purify Flag-tagged Rpc160 Pol III complexes and associated proteins. Precipitated proteins were eluted with Flag peptide, followed by SDS-PAGE and immunoblot analysis with an anti-Flag or anti-SUMO antibody. **(B)** The eluant from (A) was subjected to anti-GFP IP using GFP-Trap beads to isolate the sumoylated species from Pol III. The beads were subsequently washed with PBS containing 8M urea and 1% SDS to remove Rpc160-associated unmodified proteins, then incubated with 2x Laemmli’s buffer at 100°C to elute sumoylated proteins. The success of the IP was confirmed by anti-SUMO immunoblot. The purified materials were subjected to tryptic digestion and analyzed by mass-spectrometry. **(C)** Flag-tagged Rpc53 was IP-ed from the indicated strains using anti-Flag beads, and detected by an anti-Flag antibody (bottom). Sumoylated Rpc53 (Su-Rpc53) was detected by anti-SUMO antibody (top). An untagged *RPC53* strain was used as a negative control. **(D)** Similar experiment as in (C) showing Rpc53 sumoylation in wild type *RPC160* cells versus *rpc160-M809I* mutant cells. **(E)** Mapping Rpc53 sumoylation sites by mutagenesis analysis. *CEN* plasmids carrying wild type or mutant Flag-tagged *RPC53* were co-transformed with a *2µ SMT3* plasmid into a wild type yeast strain. Rpc53-Flag proteins were purified with anti-Flag IP, followed by SDS-PAGE and immunoblot analysis with anti-Flag (bottom) or anti-SUMO antibody (top). **(F)** An *rpc128-A704T* strain carrying a *URA3 RPC128* plasmid was transformed with *LEU2*-based *RPC128* or *RPC53* plasmids, then grown on 5FOA medium, which forces the cells to lose the *URA3 RPC 128* plasmid. *rpc53-3KR* (K51,115,236R) rescued the growth of *rpc128-A704T*, whereas N-terminal SUMO fusion *(Su-rpc53-3KR)* abolished the rescue effect. The rescue effect is dominant because all the cells in this experiment contain wild type *RPC53* in the genome. **(G)** Similar plasmid shuffle experiment as in Figure 2A. The *LEU2* plasmids carrying the indicated *RPC53* alleles were transformed into an *rpc53Δ* strain containing a *URA3 RPC53* plasmid. The transformants were then plated onto a 5FOA plate to lose the *URA3 RPC53* plasmid, and the results showed that the N-terminally SUMO-fused Rpc53 protein *(Su-rpc53-3KR)* fully supports cell viability.

Rpc53 and Rpc37 form a subcomplex in the Pol III holoenzyme (Hoffmann et al., 2015; Kassavetis, Prakash, & Shim, 2010; Landrieux et al., 2006). Identifying both of them suggests that they are either tightly associated with a sumoylated protein or are sumoylated themselves. Indeed, Rpc53 was extensively sumoylated in vivo, and this was largely dependent on Siz1 but not Siz2 (Figure 3C), correlating with the fact that *siz2A* did not rescue the *rpc160* mutant growth defect (Figure 1C). Rpc53 was sumoylated more extensively in *rpc160* mutant cells (Figure 3D), suggesting that it may serve as a better SUMO substrate when Pol III is defective. The major sumoylation sites were mapped to K51, K115, and K236, by showing that mutating all three of them to arginines *(K51, 115, 236R*, or *rpc53-3KR)* abolished the majority of sumoylation, and no modification was detected when the N-terminal 274 amino acids of Rpc53 were deleted (Δ2-275) (Figure 3E). Importantly, *rpc53-3KR* rescued the *rpc128-A704T* growth defect (Figure 3F), whereas SUMO fusion to the N-terminus of Rpc53 *(Su-rpc53-3KR)*, which mimics constitutive sumoylation, abolished the rescue effect of Rpc53-3KR. The SUMO-Rpc53-3KR fusion protein was expressed and functional, as it fully complemented the growth defect of *rpc53* null (Figure 3G). The rescue by *rpc53-3KR* was partial, suggesting additional modification sites in Rpc53 or other relevant SUMO substrates exist. Nevertheless, these results confirmed a direct relationship between SUMO and Pol III, and suggest that Rpc53 sumoylation can repress a defective Pol III machinery.

### Pol III is repressed by ubiquitylation and the Cdc48 segregase

Sumoylation itself is not sufficient to inhibit Pol III and the effect of SUMO seems to be indirect, based on the facts that constitutive sumoylation of Rpc53 did not lead to any growth defect (Figure 3G), and that *rpc160-M809I* could also be rescued by a SUMO protease mutant, *ulp2-101* (Figure S4A), which did not abolish Rpc53 sumoylation (Figure S4B). Therefore, it is likely that Rpc53 sumoylation triggers a downstream event, such as STUbL-mediated ubiquitylation, which in turn represses Pol III. Indeed, the full repression of Pol III also requires ubiquitylation, as deletion of either one of the STUbL subunits (Slx5 and Slx8), or the Ubc4 ubiquitin E2 enzyme could all rescue the growth defect caused by *rpc160* mutations (Figure 4A, 4B, and 4C). Furthermore, we could detect a physical interaction between Slx5 and Rpc53 in a yeast two-hybrid assay, which required Rpc53 sumoylation, as the 3KR mutation or the N-terminal deletion of Rpc53 abolished this interaction (Figure 4D). Consistently, the SUMO-interacting motifs (SIMs) in Slx5 are required for it to repress Pol III, as expression of the Slx5 SIM mutant did not rescue *rpc160* mutations in the *slx5A* strain (Figure 4E). These results are consistent with the STUbL activity of Slx5-Slx8 complex being important, and suggests that ubiquitylation acts downstream of sumoylation in Pol III repression.

**Figure 4.**
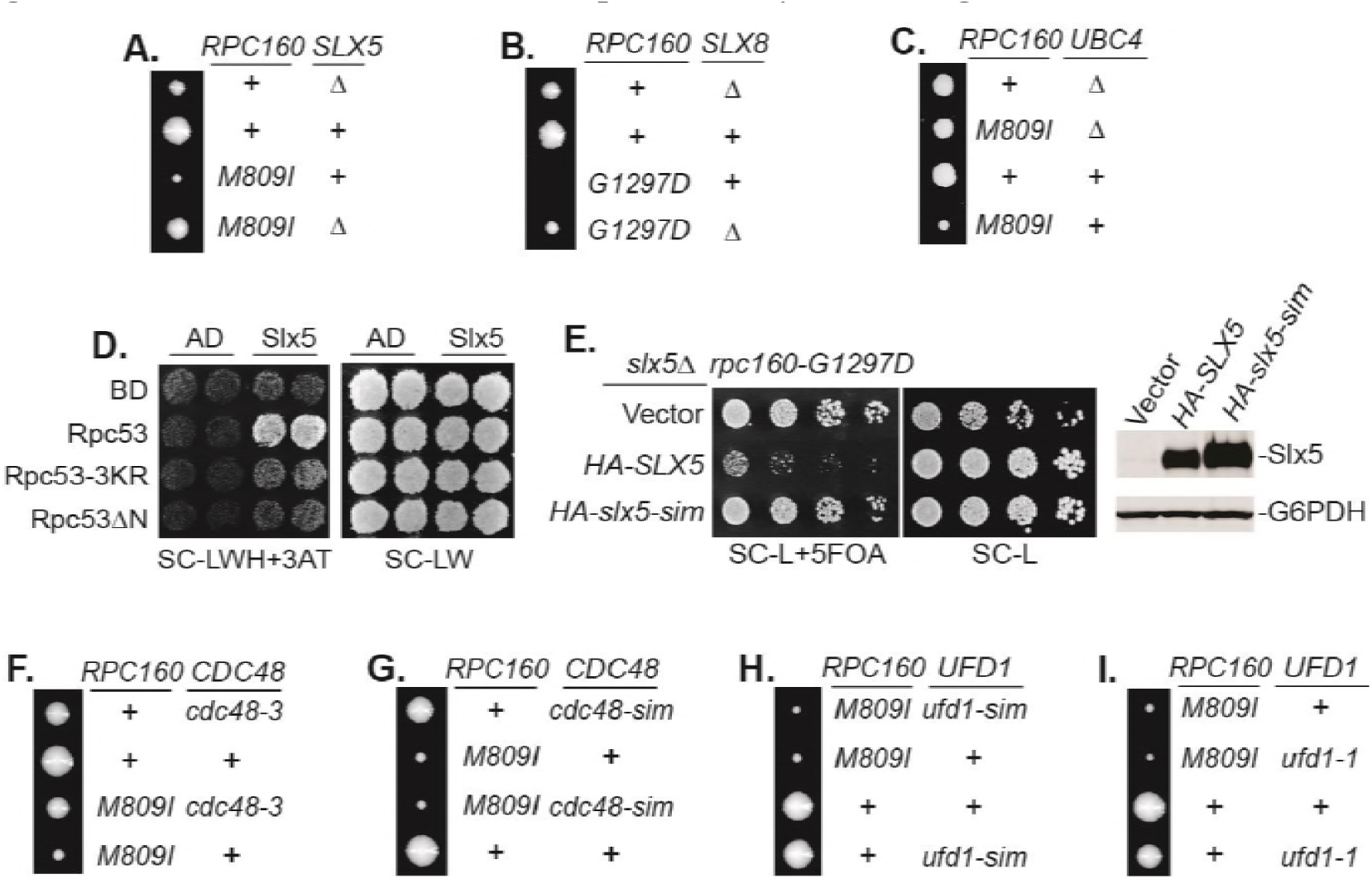
Pol III is repressed by ubiquitylation and p97/Cdc48. **(A-C)** The indicated *rpc160* mutant strains were crossed with *slx5Δ, slx8Δ*, or *ubc4Δ* strain, respectively, followed by tetrad analysis. The cross between *slx8A* and *rpc160-G1297D* was shown, because *slx8Δ* caused obvious growth defect by itself, so the rescue effect was more obvious on *rpc160-G1297D* which is a sicker mutant than *rpc160-M809I.* **(D)** Yeast two-hybrid interactions between Slx5 and Rpc53. *SLX5* and *RPC53* were cloned into a *2µLEU2* Gal4 activation-domain (AD) vector and a 2µ*TRP1* DNA-binding domain (BD) vector, respectively, and co-transformed into yeast strain PJ69-4A. Transformants were selected on synthetic media lacking leucine and tryptophan (SC-LW), then patched and replica plated to selective media lacking histidine to test for interactions. The histidine-lacking media was supplement with 3-aminotriazole (SC-LWH+3AT) for a more stringent phenotype. **(E)** *LEU2* plasmids carrying HA-tagged wild type or SIM-defective *SLX5 (HA-slx5-sim)* were transformed into an *rpc160-G1297D slx5A* strain containing a *URA3 RPC160* plasmid. Transformants were selected on SC-L then spotted onto an SC-L+5FOA plate to lose wild type *RPC160. HA-SLX5* complemented *slx5A* so the cells became sicker compared to the empty vector control transformants, while *HA-slx5-sim* did not complement, indicating that the SIMs are essential for the function *of SLX5* in this assay. The lost Slx5 function by the SIM mutations was not caused by insufficient proteins, since there were comparable levels of Slx5 proteins, as determined by an anti-HA immunoblot on total cell lysates (right, top panel). G6PDH served as a loading control (right, bottom panel). **(F-I)** Tetrad analysis between *rpc160-M809I* and *cdc48-3, cdc48-sim, ufd1-sim*, and *ufd1-1*, respectively.

Sumoylated and ubiquitylated proteins can both be targeted by Cdc48, leading us to test whether it is required in this case. As expected, *rpc160-M809I was* similarly rescued by the *cdc48-3* mutation (Figure 4F). However, the SIMs in Cdc48 (Figure 4G) or its cofactor Ufd1 (Figure 4H) were not required for its repressive effect, suggesting that Cdc48 activation does not occur through direct recognition of sumoylated Pol III complexes, but more likely through recognition of a ubiquitylated protein instead. The *ufd1-1* mutant has defects in the ubiquitin fusion degradation pathway (Johnson, Ma, Ota, & Varshavsky, 1995), but did not rescue *rpc160-M809I* (Figure 4I), suggesting Ufd1 is not the cofactor used by Cdc48 in this case.

### Pol III repression is partially mediated by ubiquitylation of Rpc160

It is conceivable that STUbL-mediated ubiquitylation represses Pol III by modifying components of the transcription machinery, including subunits of Pol III itself. We noticed that the mutant Rpc160-M809I proteins are less stable than wild type Rpc160, as determined by a cycloheximide-chase experiment (Figure 5A), suggesting the mutant Rpc160 proteins are degraded and therefore Rpc160 is likely to be ubiquitylated. Rpc160 can be ubiquitylated at 1240, K1242, K1249, K1273, and K1432, as determined by a previous proteomic study (Swaney et al., 2013). When the three clustered lysines were mutated to arginines (K1240, 1242, 1249R or 3KR), Rpc160-M809I proteins became more stable (Figure 5A) and the phenotypes of *rpc160-M809I*, including slow growth (Figure 5B) and reduced tRNA levels (Figure 5C), were partially rescued. The rescue effect was more obvious on the *rpc160-G1297D* mutant (Figure 5D). Interestingly, the 3KR mutation, when introduced into wild type Rpc160 proteins, could rescue the defect caused by mutations in a different Pol III component, such as *rpc128-A704T* (Figure 5E). In addition, the Rpc160-M809I proteins were similarly stabilized in *siz1Δ, rpc53-3KR, ubc4Δ, slx8Δ* and *cdc48-3* cells (Figure 5F), as well as in the presence of a proteasome inhibitor, MG132 (Figure 5G). These results suggest that Rpc53 sumoylation leads to Rpc160 ubiquitylation by the Slx5-Slx8 STUbL, which subsequently triggers Pol III disassembly by the Cdc48 segregase, and eventually results in Rpc160 degradation by the proteasome. The rescue by 3KR is partial, suggesting other ubiquitylation sites in Rpc160 and/or additional ubiquitylated proteins exist that play a role.

**Figure 5.**
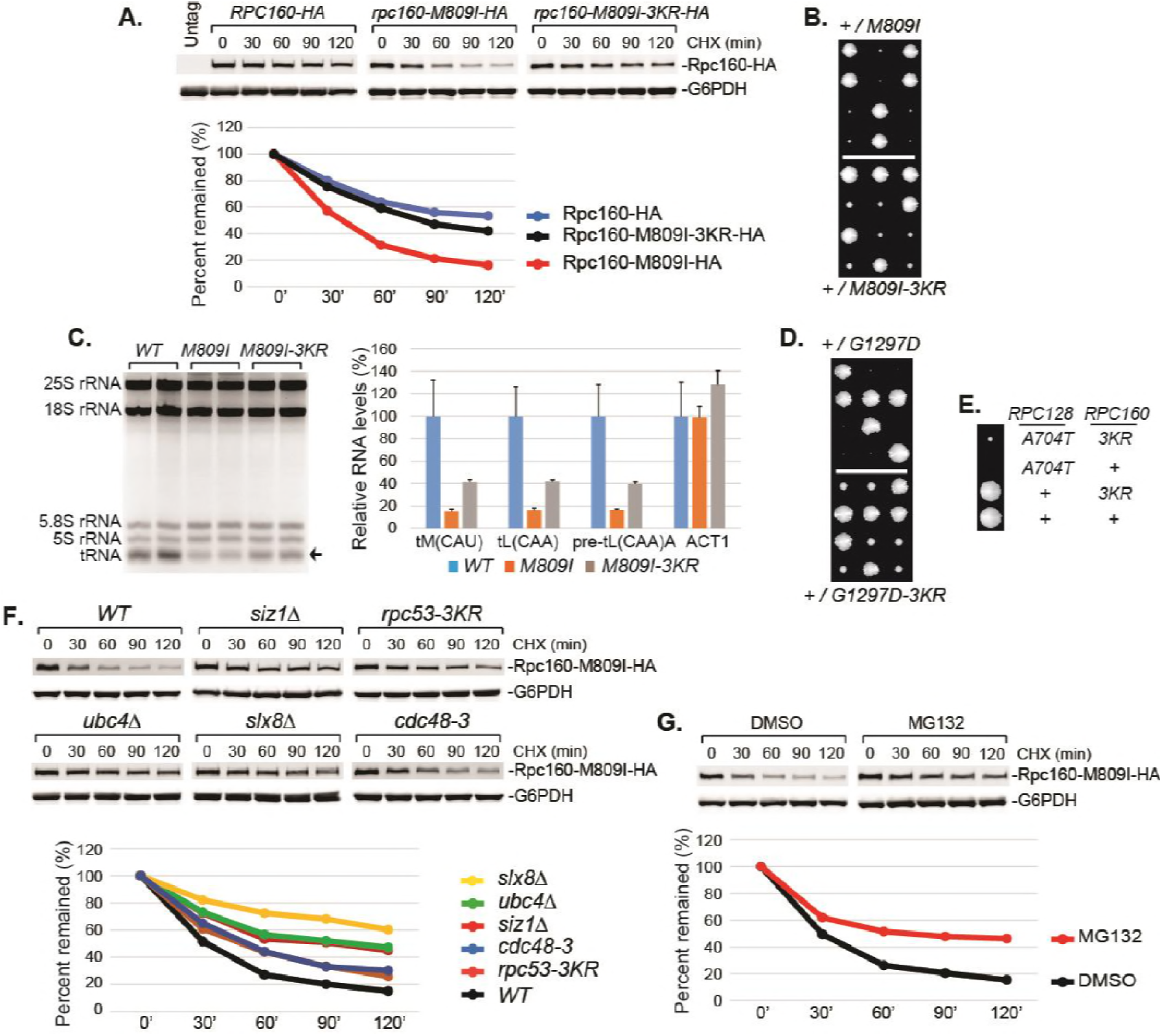
Pol III repression by ubiquitylation is partially mediated through Rpc160. **(A)** *CEN URA3* plasmids expressing HA-tagged wild type or mutant Rpc160, as indicated, were transformed into a wild type strain, and their stabilities were assayed during a cycloheximide (CHX) chase time course. Rpc160 was detected by an anti-HA antibody, and G6PDH was used as a loading control. Quantification of the bands was shown below the immunoblot. **(B)** Tetrad analysis of the diploid strains, *RPC160+/rpc160-M809I (top)* and *RPC160+/rpc160-M809I-3KR* (bottom). Tetrads from these two diploids were dissected and plated on the same YPD plate at the same time, in order to compare the growth of *rpc160-M809I* and *rpc160-M809I-3KR* cells. The growth of three dissected tetrads were shown. The large colonies are wild type *RPC160* cells, and the small colonies are *rpc160-M809I* (top) or *rpc160-M809I-3KR* (bottom) cells. The *rpc160-M809I-3KR* cells grew slightly faster than the *rpc160-M809I* cells. **(C)** Left: 2 **n**g of RNA extracted from the indicated strains was run on a 2.8% agarose gel containing ethidium bromide, then visualized with UV. Two colonies were picked for each strain. Right: RNA from left was reverse transcribed into cDNA, followed by real-time PCR analysis, as described in Figure 2E. GAPDH transcripts were used as loading control. **(D)** Similar tetrad analysis as in (B) of the diploid strains, *RPC160+/rpc160-G1297D* (top) and *RPC160+/rpc160-G1297D-3KR* (bottom). Large colonies are wild type *RPC160* cells, while the missing colonies (top) are *rpc160-G1297D* cells, and the small colonies (bottom) are *rpc160-G1297D-3KR* cells. **(E)** An *rpc128-A704T* strain was crossed with an *rpc160-3KR* strain, followed by tetrad analysis. **(F)** A *CEN URA3 rpc160-M809I-HA* plasmid was transformed into the indicated strains, and the stabilities of the Rpc160-M809I-HA proteins were determined by CHX chase time course, as described in (A). **(G)** A *CEN URA3 rpc160-M809I-HA* plasmid was transformed into a wild type strain, and protein stabilities were determined by CHX chase experiment in the presence of DMSO or MG132.

### Sumoylation and ubiquitylation of Pol III occur on the chromatin

Chromatin association of SUMO at tRNA genes was previously reported in yeast (Chymkowitch et al., 2017) and mammalian cells (Neyret-Kahn et al., 2013). It is thus possible that sumoylation and ubiquitylation of Pol III both occur on the chromatin. Indeed, we could detect an enrichment of Siz1, Slx5, as well as Cdc48-3 at tRNA genes, especially in the mutant *rpc160-M809I* cells (Figure 6A, 6B, **and** 6C). Chromatin association of wild type Cdc48 could not be detected (data not shown), possibly because it continuously disassembles the ubiquitylated Pol III complexes, releasing them as well as itself, from the chromatin. Surprisingly, Slx5 did not require its SIMs to associate with tRNA genes (Figure 6B), indicating the role of SUMO is not to recruit the STUbL to Pol III. Interestingly, however, the DNA-binding domain (SAP domain) of Siz1 (Parker et al., 2008; Reindle et al., 2006) was required for its tRNA gene association (Figure 6A). Furthermore, the SAP domain was also required for Siz1 to sumoylate Rpc53 (Figure 6D) and to inhibit the growth of the *rpc160* mutant cells (Figure 6E). Besides Siz1, the STUbL subunit Slx8 also contains a DNA-binding activity, which was mapped to the N-terminal 163 amino acids (Yang, Mullen, & Brill, 2006). Similarly, Slx8 requires this DNA-binding domain to inhibit the growth of the *rpc160* mutant cells (Figure 6F). However, the DNA-binding domain is not required for Slx8 to associated with chromatin (data not shown), as previously reported (Yang et al., 2006). Therefore, the targeting of Pol III by SUMO and ubiquitin is likely to occur on the chromatin, and require a physical interaction between DNA and the modifying enzymes.

**Figure 6.**
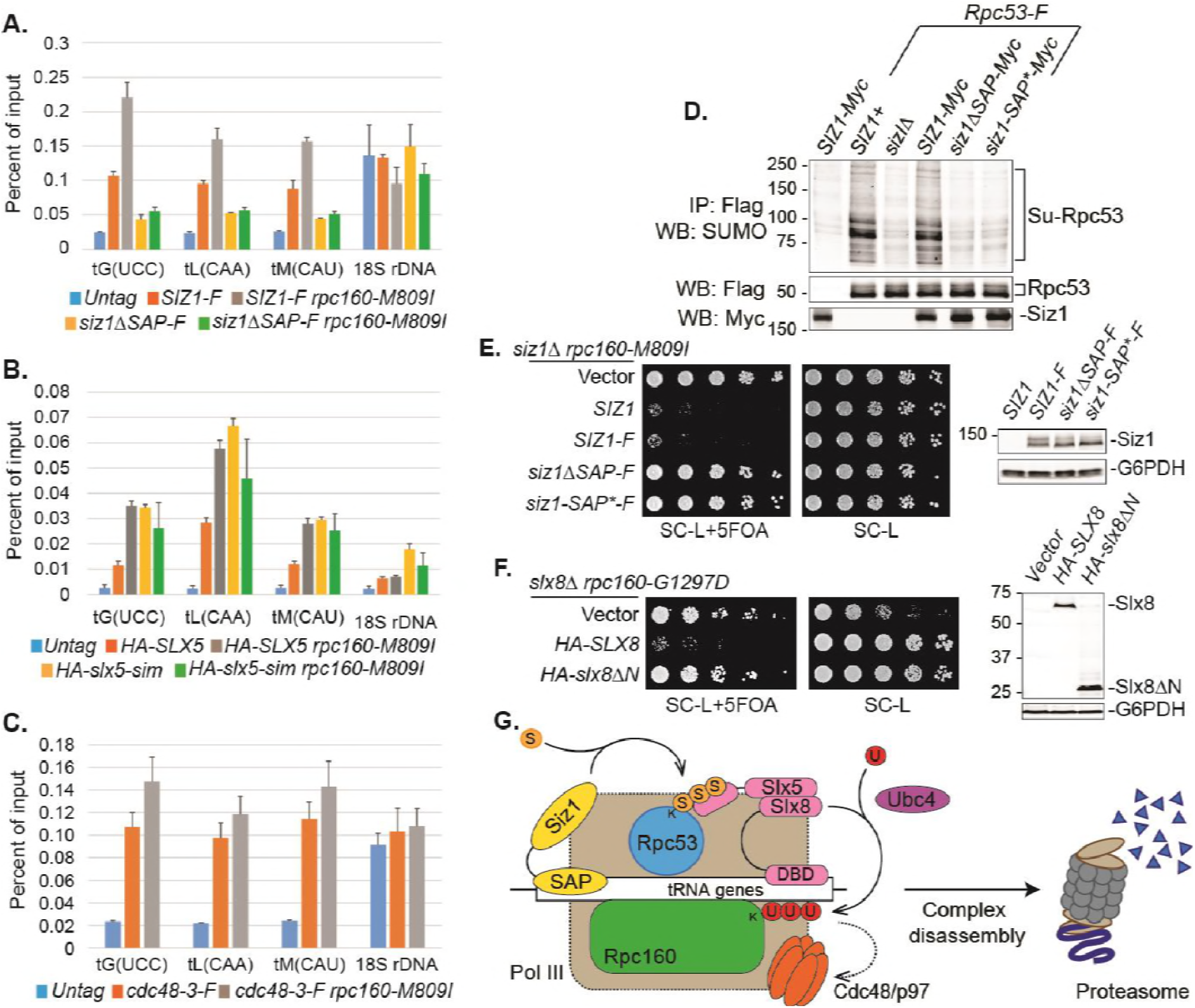
DNA is involved in Pol III repression. **(A-C)** Chromatin IP of Siz1-Flag, HA-Slx5, and Cdc48-3-Flag, respectively, using anti-Flag or anti-HA beads depending on the tag. An untagged strain was used as negative control. Chromatin association was determined by real-time PCR of the indicated genomic loci, using the percent of input method. Data are mean ± standard deviation calculated from 6 data points (two biological replicates and three technical replicates). **(D)** Flag-tagged Rpc53 proteins were purified from the indicated wild type or *siz1* mutant strains, using anti-Flag beads, followed by SDS-PAGE and immunoblotting using an anti-SUMO antibody. Rpc53 was detected by an anti-Flag antibody, and Siz1 was detected by an anti-Myc antibody. Either truncation (ΔSAP) or point mutation (SAP*) of the SAP domain resulted in loss of Rpc53 sumoylation **(E)** Left: *LEU2* plasmids carrying wild type or mutant *SIZ1* alleles were transformed into an *rpc160-M809I siz1 Δ* strain containing a *URA3 RPC160* plasmid. Transformants were selected on SC-L plate, then spotted in 5-fold dilution onto a SC-L+5FOA plate. Wild type *SIZ1* complemented *siz1 Δ* so the cells became sick on SC-L+5FOA plate, while the SAP mutants did not complement. Right: Comparable amounts of wild type and mutant Siz1 proteins were determined by anti-Flag immunoblotting on whole cell lysate, using G6PDH as loading control. **(F)** Similar plasmid shuffle experiment as in (E). *LEU2* plasmids carrying HA-tagged wild type *SLX8* or *slx8AN* (*Δ*2-163) were transformed into an *rpc160-G1297D slx8Δ* strain containing a *URA3 RPC160* plasmid. Wild type *SLX8* complemented *slx8Δ*, while *slx8ΔN did* not. Comparable amounts of Slx8 proteins were determined by an anti-HA immunoblot. **(G)** Model of Pol III regulation by SUMO, ubiquitin, and Cdc48. A stable interaction between chromatin and the SAP domain of Siz1 stimulates its activity to modify Rpc53 with SUMO (S). Rpc53 sumoylation triggers ubiquitin (U) modification of Rpc160 and potentially other proteins by the Slx5-Slx8 complex, which also required the interaction between chromatin and the DNA-binding domain (DBD) of Slx8. Ubiquitylation subsequently activates Cdc48 to disassemble the Pol III complex, facilitating degradation of Pol III subunits by the proteasome.

## DISCUSSION

Sumoylation is a very common posttranslational modification, with thousands of identified sumoylation sites in mammalian cells. Deletion of the *UBC9* SUMO-conjugating enzyme gene is an embryonic lethal event in mice underscoring the importance of sumoylation (Nacerddine et al., 2005). However very few of the known sumoylation sites have been shown to have a functional consequence, because mutation of single sumoylation sites or even combinations in a protein usually results in no obvious phenotype. A possible explanation is provided by the “SUMO spray” model, which proposes that a locally concentrated SUMO E3 ligase sumoylates multiple proteins nearby, allowing SUMO to serve as a glue for protein complex assembly (Psakhye & Jentsch, 2012). In such cases, sumoylation on multiple proteins would need to be abolished simultaneously to reveal a phenotype, making genetic analysis of SUMO function more difficult. To address this challenge, we devised, a phenotype-based genetic screen that selects for point mutations in yeast whose growth is rescued only when sumoylation is compromised, allowing identification of proteins where sumoylation has a functional consequence that can then be studied.

### SUMO-ubiquitin-Cdc48 is a new regulatory pathway for Pol III

We used a yeast genetic approach to uncover a functional relationship between Pol III and SUMO, demonstrating that genetics is a powerful tool to study sumoylation, complementary to the biochemical approach. Performing the same type of screen under different conditions is likely to yield more functional SUMO targets in the cell, and the same principle can be potentially extended to study other posttranslational forms. Our findings support a model where SUMO, ubiquitin, and Cdc48 act in a linear pathway to repress Pol III transcription (Figure 6G). In this model, a defective/stalled chromatin-associated Pol III complex on a tRNA gene is first recognized by the chromatin-associated Siz1 E3 SUMO ligase and sumoylated on the Rpc53 subunit. Rpc53 sumoylation would then trigger ubiquitylation of the Rpc160 Pol III catalytic subunit, and possibly other proteins by the chromatin-associated Slx5-Slx8 SUMO-targeted E3 ubiquitin ligase complex. Subsequently, ubiquitylated Pol III complexes are recognized and disassembled by the Cdc48 AAA-ATPase segregase, leading to proteasomal degradation of the Rpc160 subunit, thus clearing obstructed tRNA genes to allow transcription to resume. We showed that this pathway is independent of the Mck1 and Kns1 kinases and Maf1, thus representing a new regulatory mechanism for Pol III. Interestingly, SUMO was recently shown to promote Pol III assembly and activity by modifying another subunit, Rpc82 (Chymkowitch et al., 2017), suggesting that SUMO modification has complex regulatory effects on Pol III. By modifying different components of the Pol III machinery, SUMO may regulate Pol III at multiple transcription steps or in response to various signaling events.

### The SUMO-ubiquitin-Cdc48 pathway may serve as a quality control mechanism for Pol III

In the particular case reported here, the SUMO-ubiquitin-Cdc48 pathway seems to preferentially target defective Pol III, based on the results that *siz1A* only has an obvious effect when the function of Pol III is impaired, that more Siz1, Slx5, and Cdc48 proteins are associated with tRNA genes, and that Rpc53 is more extensively sumoylated in mutant *rpc160* cells. The question is, what Pol III defect is being recognized by the pathway? Since mutations in the Pol III initiation factors Brf1, Tfc1, and Tfc6 can also be rescued by disrupting sumoylation, it is likely that a defect in transcription initiation is being recognized. This is supported by the result that *siz1 Δ* rescued an initiation-defective mutant, *rpc31-236.* In Rpc160, the mutations T379I (close to the catalytic site), A880T (in bridge helix), and G1098T (in trigger loop) (Hoffmann et al., 2015) are likely to impair elongation, suggesting elongation defects might also be a feature recognized by the SUMO-ubiquitin-Cdc48 pathway. Consistently, two previously characterized elongation mutants, *rpc 160–112* and *rpc160-270*, were both rescued by *siz1 Δ.*

The next question is, how are initation- and elongation-defective Pol III proteins targeted by the pathway? Our data imply that it is the whole Pol III complex rather than individual subunits that are targeted for repression. For instance, sumoylated Rpc53 was co-purified with Rpc160, suggesting it exists in the complex. In addition, when a mutation occurs in one subunit (e.g. Rpc128), SUMO and ubiquitin are conjugated to different subunits (Rpc53 and Rpc160, respectively), instead of Rpc 128 itself, and the mutant Rpc 128 proteins did not become unstable (data not shown). It is possible that SUMO and ubiquitin recognize overall conformational changes in the Pol III protein complex caused by a defective subunit, but we favor another possibility that involves chromatin DNA. It is conceivable that initiation or elongation defects will trap Pol III on the chromatin, forming a relatively stable protein/DNA complex. It would be a more efficient way to distinguish defective Pol III from normal Pol III molecules by utilizing the stable interaction between the E3 ligases and chromatin DNA, rather than by recognizing conformational changes. In fact, Siz1, Slx5, and Cdc48 are all associated with tRNA genes, and both Siz1 and Slx8 contain DNA-binding domains that are required for them to repress Pol III. The requirement of the DNA-binding activities in Siz1 and Slx8 is somewhat surprising, since no functional consequences have been reported when they are disrupted. Specifically, the SAP-truncated version of Siz1 can still sumoylate PCNA and maintain the DNA-damage sensitivity of the *rad18* mutant, which can be reversed by complete deletion of *SIZ1* (Parker et al., 2008). Similarly, deleting the N-terminal DNA-binding domain of Slx8 does not affect its ability to associate with chromatin (Yang et al., 2006), or to complement the *slx8A* synthetic lethal phenotype with *sgs1Δ.* Therefore, the requirement for DNA-binding activity may indicate a role of SUMO and ubiquitin in a process other than the DNA damage response or genome stability maintenance.

Unlike the SAP domain in Siz1, the SIMs in Slx5 and the DNA-binding domain in Slx8 are not required for their chromatin association at tRNA genes, suggesting that their main function is not to recruit STUbL to Pol III. Instead, they may provide important docking sites to position the enzyme in the right orientation relative to the substrate in order for it to ubiquitylate a specific target subunit in Pol III. It is thus possible that the STUbL can travel with sumoylated Pol III without ubiquitylating the polymerase until Pol III is somehow trapped on the chromatin, which will allow STUbL to stably bind to DNA and activate its ligase activity. Therefore, Rpc53 sumoylation itself will not be sufficient to trigger ubiquitylation to inhibit Pol III. This is supported by the finding that SUMO-fused Rpc53 did not affect cellular growth (Figure 3G), even though the N-terminal SUMO fusion could functionally replace sumoylation on the natural modification sites (K51, K115, and K236) (Figure 3F). These results also suggest that sumoylation may not activate Cdc48 directly, but rather indirectly through ubiquitylation. Consistently, the SUMO-interacting activities of Cdc48 and its cofactor Ufd1 were not required for Pol III repression, although how Cdc48 is activated and what cofactors are required for Cdc48 in this case remain to be answered.

Taken together, we propose that SUMO, ubiquitin, and Cdc48 act in a sequential manner, and that together with the additional requirement of DNA-binding activities, they confer substrate specificity to restrict ubiquitylation, as well as subsequent complex disassembly and proteasomal degradation, only towards transcriptionally defective Pol III, while leaving normal Pol III unaffected. To further test our hypothesis, in vitro sumoylation, ubiquitylation, and Pol III transcription assays with purified proteins are required.

### The SUMO-ubiquitin-Cdc48 pathway is a potential target for Pol III-related human diseases

Given the fact that Pol III, sumoylation, ubiquitylation, and Cdc48 are all conserved from yeast to humans, this new Pol III regulatory mechanism is likely to be conserved as well. In fact, proteomic studies of sumoylation in mammalian cells have identified sumoylated proteins in Pol III, including four Pol III-specific subunits and two subunits shared by Pol I and/or Pol II (Table S4) (Hendriks et al., 2014; Lamoliatte et al., 2014; Tammsalu et al., 2014). By comparison, only three subunits for each of Pol I and Pol II were found to be sumoylated, suggesting Pol III is the major SUMO target among the three polymerases. This correlates with the specific genetic relationship between SUMO and Pol III observed in yeast, namely that *siz1Δ* only rescued mutations in Pol III but not those in Pol I or Pol II. Interestingly, RPC4, the human homologue of yeast Rpc53, seems to be the most prevalent sumoylated protein among all human RNA polymerase subunits, with a total number of 10 sumoylation sites combining all three datasets. Mutations in Pol III cause neurodegenerative disorders in humans (Bernard et al., 2011; Borck et al., 2015; Saitsu et al., 2011; Shimojima et al., 2014; Synofzik et al., 2013; Terao et al., 2012; Tetreault et al., 2011; Thiffault et al., 2015). We showed, interestingly, that the phenotypes caused by a subset of these Pol III disease mutations, when introduced into yeast cells, were rescued when sumoylation was disrupted (Figure S3 **and** Table S3). In addition, the fibroblasts derived from a patient carrying Pol III mutations exhibited reduced levels of tRNA (data not shown), as is the case in yeast. It is thus intriguing to speculate that sumoylation, ubiquitylation, and Cdc48 segregase can all be potential therapeutic targets for the neurodegenerative disease caused by Pol III mutations. The next step is to determine if this regulatory mechanism is conserved in humans, by testing it in cultured mammalian cells and mouse model systems. We have shown that the yeast growth assay is a convenient tool to determine which neurodegenerative disease mutations are likely to cause a phenotype and therefore, which should be chosen to create cell lines and mouse models. It is possible to differentiate iPSC lines into myelinating oligodendrocytes in vitro (Kerman et al., 2015), and the oligodendrocytes from Pol III patients are expected to display a myelination defect. Alternatively, a cell line can be made to carry a lethal mutation in one copy of a Pol III subunit gene while leaving the wild type copy under control of an inducible promoter. Such a cell line can be adapted for high-throughput chemical screens for inhibitors against sumoylation, ubiquitylation, or Cdc48, which can not only be used as research tools, but also be developed into potential therapies for Pol III-related disorders or other human diseases involving SUMO, ubiquitin, or Cdc48, such as cancer (Kessler et al., 2012). A genetically modified mouse model will eventually be needed to recapitulate the disease and test the effect of the inhibitors.

## Experimental Procedures

### Yeast strains, plasmids, media, and genetic methods

The *Saccharomyces cerevisiae* strains and plasmids used in this study are listed in Table S5 **and** Table S6, respectively. All media used, including rich YPD medium (yeast extract-peptone-dextrose), sucrose medium (yeast extract-peptone-Suc), synthetic complete (SC) drop-out medium (for example, SC-U), SC-galactose medium and sporulation medium, were made as described previously (Rose, 1990). SC-L+5FOA plates were made as standard SC drop-out medium, but using 2 g of SC-UL drop-out mix, plus 50 mg of uracil, and 1 g of 5FOA per 1 liter of total volume. For a more stringent yeast two-hybrid interaction signal, 24 µmol of 3-aminotriazole (3AT) was spread onto a 10 cm SC-LWH plate. Standard genetic methods for mating, sporulation, transformation, and tetrad analysis were used throughout this study. In the tetrad analysis experiments, the mutant haploid *rpc160* or *rpc128* strains contain a *URA3* vector carrying wild type copy of gene, in order to maintain the strains. The *URA3* plasmids were lost from the diploid cells on 5FOA media after mating and before sporulation. The similar strategy was used in the plasmid shuffle experiments, for example, in Figure 2A, where the starting *rpc160Δ* strain contains a *URA3 RPC160* plasmid. Upon transformation with *LEU2* plasmids carrying *rpc160* mutant alleles, the transformants were plated onto 5FOA-containing media to lose the *URA3 RPC160* plasmid. Growth on 5FOA media therefore reflects the growth phenotype of the *rpc160* mutants present on the *LEU2* plasmids.

The mutations in *slx5-sim* are: _24_VILI – VAAA, _93_ITII – ATAA, _116_YVDL – YAAA, and _155_LTIV – ATAA. The *siz1ΔSAP* and *siz1-SAP\** mutations were made as previously described (Parker et al., 2008).

### Design of the reverse suppressor screen of *SMT3-Q56K.*

The starting strain is ZOY261 *(ade2 ade3 ura3 leu2 trp1 can1A::FUR1::natMX4)* carrying two plasmids, pZW321 *(CEN URA3 ADE3 SMT3-Q56K)* and pAK12-1 *(CEN TRP1 ade3-pink)* (Koren, Ben-Aroya, Steinlauf, & Kupiec, 2003), and grown on SC-Leu-Trp in order to keep the plasmids. Wild type yeast cells are white, while *ade2* mutant is red. *ade3* mutation suppresses *ade2*, so that *ade2 ade3* double mutant is white. Both *ade2* and *ade3* are recessive, so in the presence of pZW321, ZOY261 colonies are red. The starting strain (ZOY261 + pZW321 + pAK12-1), if grown on YPD, does not need pZW321 for viability, so the cells will lose the plasmid during cell proliferation, eventually forming colonies with red and white sectors. The screen looks for mutations that cause sickness or lethality, but can be suppressed / rescued by *SMT3-Q56K* on pZW321. These mutants will appear as uniformly red colonies, because they always need to keep pZW321 for viability. They will also be sensitive to 5FOA, because 5FOA counter-selects against *URA3* on pZW321.

To perform the screen, cells were mutagenized with 3% ethyl methanesulfonate (EMS), washed, and then spread onto YPD plates at 30°C to allow formation of single colonies. Uniformly red colonies were first picked and restreaked on fresh YPD plates. The clones remaining uniformly red after restreak were subsequently screened for those that are 5FOA-sensitive. The red 5FOA-sensitive colonies could also come from mutations that are synthetic lethal with the *ura3* or *ade3* alleles, as pZW321 also carries wild type *URA3* and *ADE3.* To reduce the chance of isolating these undesired mutations, two modifications were made. First, an additional copy of wild type *FUR1* gene was integrated at the *CAN1* gene locus of the starting strain, because *ura3* synthetic lethal mutations are most frequently found in *FUR1* (Koren et al., 2003). Second, the pAK12-1 plasmid carrying an *ade3-pink* allele was co-transformed with pZW321 into ZOY261. Unlike wild type *ADE3*, the *ade3-pink* allele is partially functional, but confers a pink (instead of red) colony color phenotype in *ade2 ade3* background (Koren et al., 2003). Therefore, in the presence of pAK12-1, the *ade3* synthetic lethal mutants will not appear as uniformly red colonies, but sectored with red and pink instead. To finally confirm that a strain contains a *SMT3-Q56K*-rescuable mutation, the strain was transformed with the pZW311 plasmid (*CEN LEU2 SMT3*-*Q56K*), which should render the cells resistant to 5FOA after transformation.

To identify the mutated gene, a wild type genomic DNA library (Jones et al., 2008) was transformed into the candidate strains, and screened for 5FOA-resistant transformants. The plasmids were then isolated and sequenced to identify the ends of the genomic DNA on the plasmids. The mutated genes were identified by subcloning or by complementation experiments. The genomic mutations were finally confirmed by PCR and sequencing. To summarize, ~80,000 colonies were initially screened, and 740 uniformly red ones were picked and restreaked. 105 clones remained red after restreak, among which 77 were 5FOA-sensitive. Finally, 25 of these 77 clones were confirmed to have mutations that can be suppressed by *SMT3-Q56K* (Table 1).

### Preparation of RNA from yeast cells

RNA was prepared by the “Heat/Freeze” method as previously described with modifications (Schmitt, Brown, & Trumpower, 1990). Briefly, yeast cells were resuspended in AE buffer (50 mM NaOAc pH 5.2, 10 mM EDTA, 1% SDS), mixed with equal volume of phenol (pH 4.5), then incubated at 65°C for 4 min. The cell suspension was then frozen on dry ice/ethanol bath and thawed at 37°C. After centrifugation at top speed, the RNA containing upper layer was transferred to a new tube. RNA was extracted first with phenol/chloroform/isoamyl alcohol (25:24:1), then with chloroform/isoamyl alcohol (24:1), and finally precipitated with 100% ethanol containing 0.3M NaOAc (pH 5.2). The RNA pellet was washed once with 70% ethanol and once with 100% ethanol, then dissolved in DEPC-treated H2O at 50°C for 10 min.

### Reverse transcription and real-time PCR analysis

RNA was converted into DNA using the SuperScript III RT kit (Invitrogen, catalog # 18080–051) with modifications. First, a mixture of random hexamer and tRNA gene-specific primers (Table S7) was used for reverse transcription. Second, primers were hybridized to RNA by incubating the sample at 100°C for 5 min, followed by 65°C for 5 min, then held at 55°C. Third, the RT enzyme mix was pre-warmed to 55°C before adding to the RNA/primer mix. Fourth, reverse transcription was carried out at 55°C for 30 min, followed by 85°C for 5 min, and finally held at 4°C. The resulting DNA was diluted 10 times with H2O, and 2 µl of the diluted DNA was used for real-time PCR. Real-time PCR was performed with SYBR Green master mix (Applied Biosystems, catalog # 4367659) on the Applied Biosystems 7900HT Fast Real-Time PCR System. Data were analyzed by the comparative Ct method (Schmittgen & Livak, 2008), using GAPDH as internal control. Primer sequences were listed in Table S6. Data are mean ± standard deviation calculated from 6 data points (two biological replicates and three technical replicates), presented as relative amount compared to wild type.

### Preparation of protein extracts and immunoprecipitation (IP)

For IPs, crude protein extracts were prepared by the glass bead beating method, as described above (Z. Wang & Prelich, 2009). Briefly, cells were first resuspended in lysis buffer containing 50 mM Tris-Cl (pH 8.0), 10 mM MgCl2, 1 mM EDTA, 150 mM NaCl, 1 mM phenylmethylsulfonyl fluoride (PMSF), 50 mM*N*-ethylmaleimide (NEM), 1% Triton X-100, and protease inhibitors (Roche, catalog # 11836170001), and glass beads. Cell lysis was subsequently performed on a Precellys 24 tissue homogenizer (setting 6500, 30 sec for 3 times), followed by centrifugation to clear the lysate. IP was performed with anti-HA agarose (Sigma, catalog #A2095) or anti-Flag-agarose (Sigma, catalog # A2220) at 4°C for a typical length of 2 hr, and the bound proteins were eluted with 2x HA or 2x Flag peptide, respectively.

For the CHX chase experiments in Figure 5 and the immunoblotting in Figure 6E **and** 6E, protein extracts were prepared by the post-alkaline extraction method, as previously described (Zhang et al., 2011). Briefly, 1–2 O.D. of cells were first washed with 1 ml of water, then resuspended in 200 µl 2 M LiOAc and incubated on ice for 5 min. The cells were subsequently resuspended in 200 µl 0.4 M NaOH and incubated on ice for another 5 min, before finally being resuspended in 40 µl Laemmli’s buffer and incubated at 100°C for 5 min. 10 µl of protein sample were loaded on the gel.

### Assays of protein stability

Yeast cultures were grown overnight at 30°C to log phase. Cell concentrations were then adjusted to OD600 = 1. To start the chase, 50 mg/ml cycloheximide (Sigma, #C7698) was added to a final concentration of 0.5 mg/ml. 1.5 ml culture was collected immediately as time point 0 in an Eppendorf tube pre-loaded with 15 µl 10% sodium azide. Cells were then washed with 1 ml water and frozen on dry ice. The remaining cultures were incubated at 30°C, and 1.5 ml samples were collected every 30 min in the same way. Crude extracts were prepared by the post-alkaline extraction method as described above. An anti-G6PDH (Sigma, #A9521) antibody was used to detect G6PDH as loading control. Immunoblot analysis was performed using the Odyssey infrared imaging system *(LI-COR* Biosciences).

### 2-step immunoprecipitation to isolate Pol III-associated sumoylated proteins

Flag-tagged Rpc160 was first purified by incubating with anti-Flag M2 affinity gel (Sigma, catalog # A2220) in lysis buffer, then eluted with 450 ng/ul 2x Flag peptide in GFP-IP buffer (50 mM Tris-Cl (pH 7.5), 1 mM EDTA, 500 mM NaCl, 1 mM PMSF, 50 mM NEM, and protease inhibitors). For the second step, the Flag-eluted protein sample was incubated with GFP-Trap agarose beads (ChromoTek, catalog # gta-10), followed by two washes with GFP-IP buffer, one quick wash with PBS containing 8 M urea and 1% SDS, and one with PBS containing 1% SDS. GFP-tagged proteins were eventually eluted with 2x Laemmli’s buffer (without dye) at 100°C for 5 min, and analyzed by mass-spectrometry. Immunoprecipitated samples were analyzed by SDS-PAGE, followed by immunoblotting with an anti-SUMO antibody (Santa Cruz, Smt3 (y-84), catalog # sc-28649) or an anti-Flag M2 antibody (Sigma). The *SLX5* gene was deleted from the strains in Figure 3A, in order to increase general sumoylation signal (Z. Wang et al., 2006; Z. Wang & Prelich, 2009).

### Mass spectrometry analysis

Samples were first denatured in 8 M urea and then reduced and alkylated with 10 mM Tris (2-carboxyethyl) phosphine hydrochloride [Roche Applied Science] and 55 mM iodoacetamide [Sigma-Aldrich] respectively. Samples were then digested over-night with trypsin [Promega] according to the manufacturer’s specifications. The protein digests were pressure-loaded onto 250 micron i.d. fused silica capillary [Polymicro Technologies] columns with a Kasil frit packed with 3 cm of 5 micron C18 resin [Phenomenex]. After desalting, each loading column was connected to a 100 micron i.d. fused silica capillary [Polymicro Technologies] analytical column with a 5 micron pulled-tip, packed with 12 cm of 5 micron C18 resin [Phenomenex].

Each split column was placed in line with an 1100 quaternary HPLC pump [Agilent Technologies] and the eluted peptides were electrosprayed directly into an Orbitrap Elite mass spectrometer [Thermo Scientific]. The buffer solutions used were 5% acetonitrile/0.1% formic acid (buffer A) and 80% acetonitrile/0.1% formic acid (buffer B). The 120 min elution gradient had the following profile: 10% buffer B beginning at 10 min to 45% buffer B at 90 min, and then 100% buffer B at 100 min continuing to 110 min. A cycle consisted of one full scan mass spectrum (300–1600 m/z) in the Orbitrap at 120,000 resolution followed by 15 data-dependent collision induced dissociation (CID) MS/MS spectra in the ion trap. Charge state screening was enabled and unassigned charge states and charge state 1 were rejected. Dynamic exclusion was enabled with a repeat count of 1, a repeat duration of 30 sec, an exclusion list size of 500 and an exclusion duration of 120 sec. Dynamic exclusion early expiration was enabled with an expiration count of 3 and an expiration signal-to-noise ratio of 3. Application of mass spectrometer scan functions and HPLC solvent gradients were controlled by the Xcalibur data system [Thermo Scientific].

MS/MS spectra were extracted using RawXtract (version 1.9.9.2) (McDonald et al., 2004). MS/MS spectra were searched with the ProLuCID (version 1.3.5) algorithm (Xu et al., 2015) against a Saccharomyces Genome Database (SGD) protein database downloaded on 01–05-2010 that had been concatenated to a decoy database in which the sequence for each entry in the original database was reversed (Peng, Elias, Thoreen, Licklider, & Gygi, 2003). A total of 13,434 protein entries were searched. Precursor mass tolerance was 50 ppm and fragment mass tolerance was 600 ppm. For protein identifications, the ProLuCID search was performed using no enzyme specificity and static modification of cysteine due to carboxyamidomethylation (57.02146). ProLuCID search results were assembled and filtered using the DTASelect (version 2.1.3) algorithm (Tabb, McDonald, & Yates, 2002), requiring full enzyme specificity (cleavage C-terminal to Arg or Lys residue) and a minimum of one peptide per protein identification. The number of missed cleavages was not specified. The protein identification false positive rate was kept below one percent and all peptide-spectra matches had less than 10 ppm mass error. DTASelect assesses the validity of peptide-spectra matches using the cross-correlation score (XCorr) and normalized difference in cross-correlation scores (deltaCN). The search results are grouped by charge state and tryptic status and each sub-group is analyzed by discriminant analysis based on a non-parametric fit of the distribution of forward and reversed matches.

### Chromatin immunoprecipitation (ChIP) and real-time PCR analysis

ChIPs were performed as previously described (Keogh, Podolny, & Buratowski, 2003). Yeast cells were grown over night at 30°C to log phase. Formaldehyde was added to a final concentration of 1% for 20 min at room temperature, and the reaction was quenched by the addition of glycine to 0.3 M. Cells were washed twice with Tris-buffered saline, and lysed with glass beads in FA lysis buffer containing 50 mM Tris-Cl (pH 8.0), 10 mM MgCl2, 1 mM EDTA, 150 mM NaCl, 1 mM phenylmethylsulfonyl fluoride (PMSF), 50 mM *N*-ethylmaleimide (NEM), 1% Triton X-100, 0.1% SDS, 0.1% sodium deoxycholate, and protease inhibitors (Roche, catalog # 11836170001). Chromatin was sheared by sonication until the average fragment size was between 200 and 500 bp.

Immunoprecipitation was performed over night at 4°C with anti-HA or anti-Flag agarose. The bound materials were eluted with 2x HA or 2x Flag peptide, followed by RNase A and protease K treatment to de-crosslink. Bound DNA was then purified with the Qiagen PCR purification kit, and analyzed by real-time PCR, as described above. Data are mean ± standard deviation calculated from 6 data points (two biological replicates and three technical replicates), using the percent of input method. An untagged strain was used as negative control. The sequences of the primers were listed in Table S7.

## AUTHOR CONTRIBUTIONS

Z.W. and T.H designed the experiments. C.W. performed the reversed suppressor screen. A.A. and J.R.Y performed M/S analysis. Z.W. and T.H. wrote the manuscript. T.H. supervised the study.

## ACKNOWLEDGEMENTS

We thank Gregory Prelich for the SUMO pathway mutant plasimds and strains and the wild type genomic DNA library, Martin Kupiec for the *ade3-pink* plasmid, Michel Werner for the *rpc160* and *rpc31* mutant plasmids, Vicki Lundblad for sharing yeast vectors and equipment, and Jill Meisenhelder for the 2x Flag peptide. We also thank Benoit Coulombe and Genevieve Bernard for sharing unpublished data on Pol III-related diseases, as well as Ian Willis, Alessandro Vannini, Gregory Prelich, and members of the Hunter lab for constructive discussions. This study was supported by NIH grants #CA080100, CA082683 and CA014195 to T.H., NCRR grant #5P41RR011823-17 and NIGMS grant #8P41GM103533-17 to J.R.Y., the Helmsley Center for Genomic Medicine, and the Functional Genomics Core at the Salk Institute. T.H. holds the Renato Dulbecco Chair in Cancer Research, and is a Frank and Else Schilling American Cancer Society Professor.

## COMPETING INTERESTS

The authors declare no competing interests.

## SUPPLEMENTARY FIGURES AND TABLES

**Table S1.**
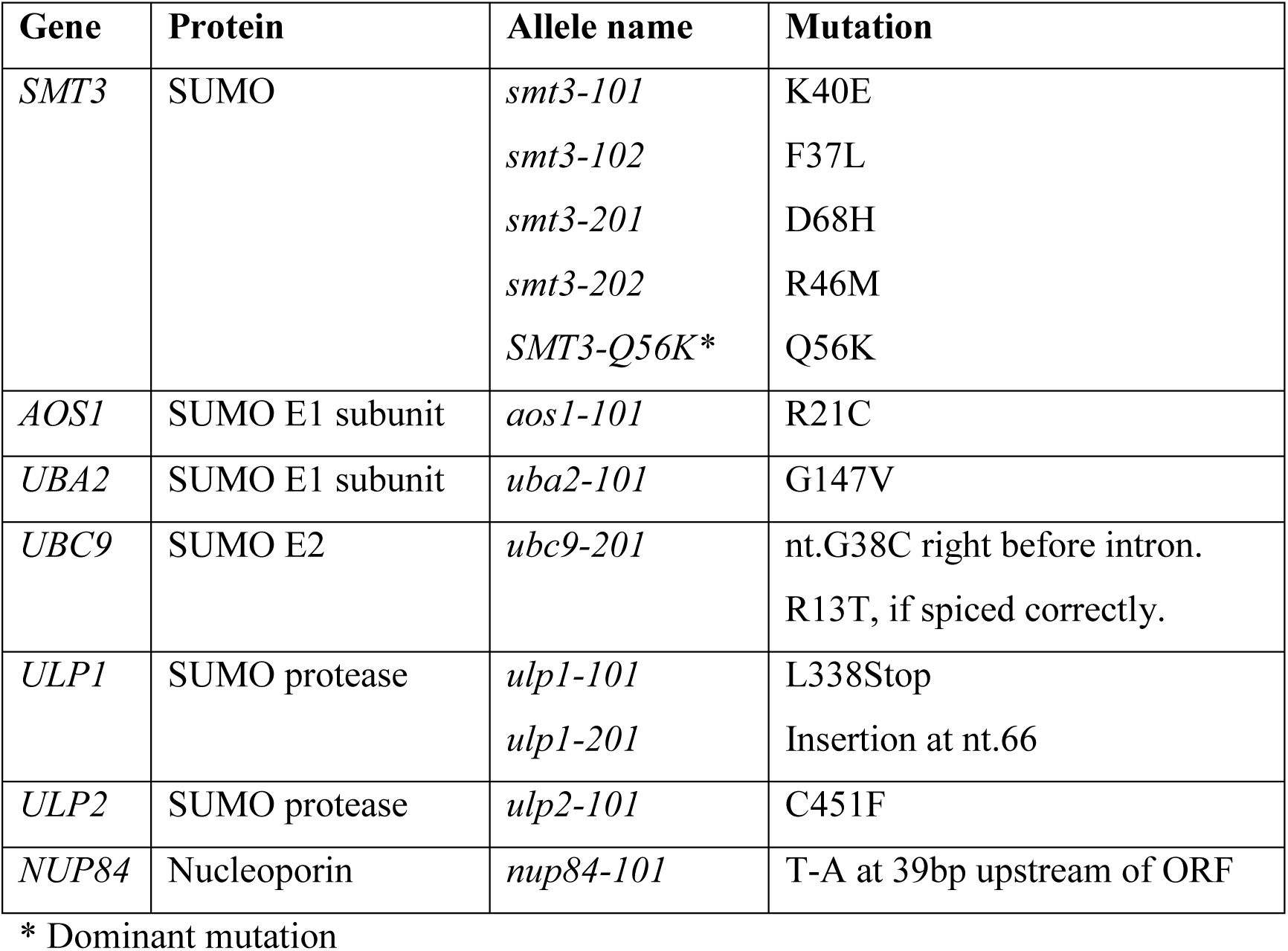
*mot1-301* suppressor mutations.

| Gene | Protein | Allele name | Mutation |
| --- | --- | --- | --- |
| *SMT3* | SUMO | *smt3-101* | K40E |
|  |  | *smt3-102* | F37L |
|  |  | *smt3-201* | D68H |
|  |  | *smt3-202* | R46M |
|  |  | *SMT3-Q56K\** | Q56K |
| *AOS1* | SUMO E1 subunit | *aos1-101* | R21C |
| *UBA2* | SUMO E1 subunit | *uba2-101* | G147V |
| *UBC9* | SUMO E2 | *ubc9-201* | nt.G38C right before intron.<br>R13T, if spiced correctly. |
| *ULP1* | SUMO protease | *ulp1-101* | L338Stop |
|  |  | *ulp1-201* | Insertion at nt.66 |
| *ULP2* | SUMO protease | *ulp2-101* | C451F |
| *NUP84* | Nucleoporin | *nup84-101* | T-A at 39bp upstream of ORF |

\* Dominant mutation

**Figure S1.**
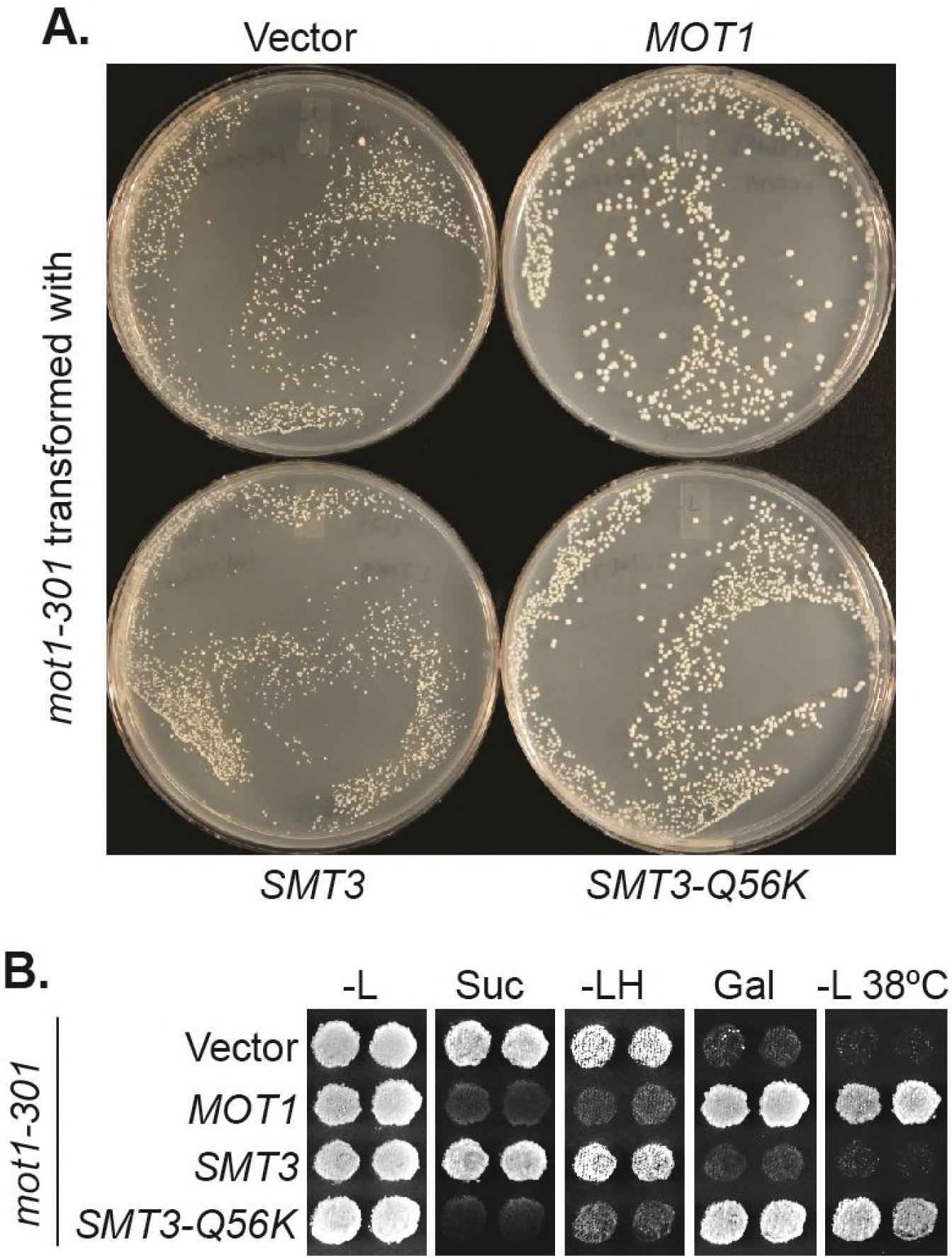
*SMT3-Q56K* suppresses *mot1-301* dominantly. **(A)** A *mot1-301* strain was transformed with *CEN LEU2* vectors carrying indicated genes, then selected for transformants on SC-Leu plates. Wild type *MOT1* or *SMT3-Q56K* made *mot1-301* cells grow faster. **(B)** Transformants from (A) were patched on SC-Leu (-L) then replica plated to SC-Leu-His (-LH), YPSucrose (Suc), SC-Galactose (Gal) plates, or a SC-Leu plate incubated at elevated temperature 38°C (-L 38°C). *mot1-301* is His^+^ (Spt^-^), Suc^+^ (Bur^-^), Gal^-^, and Ts^-^, whereas *SMT3-Q56K* reversed all four phenotype. The suppression is dominant because the wild type genomic copy of *SMT3* was present in all the strains.

**Figure S2.**
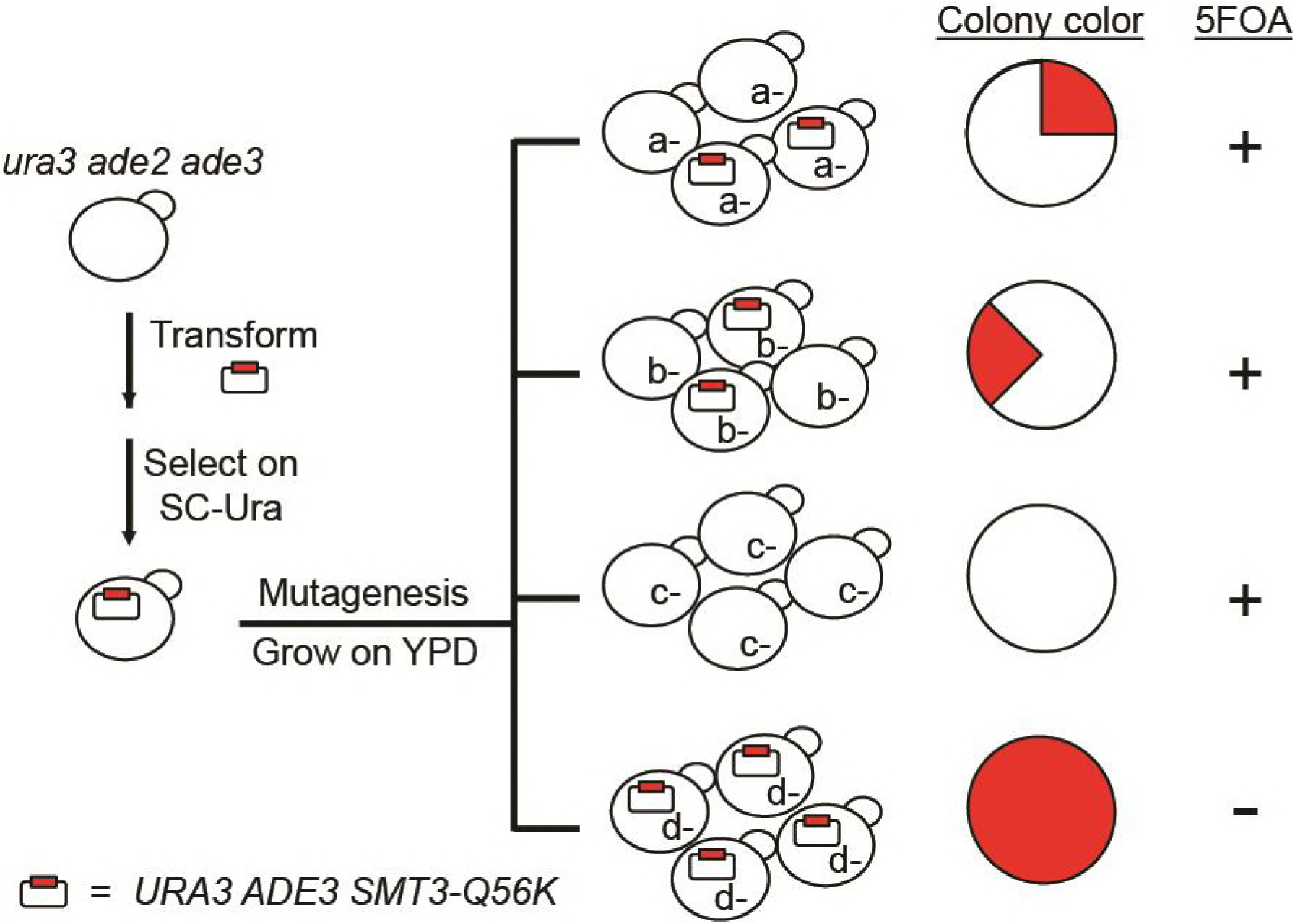
Design of the SUMO reverse suppressor screen. The starting strain is an *ura3 ade2 ade3* triple mutant. *ura3* is used for *URA3* plasmid selection and 5FOA-sensitivity test. *ade2 ade3* double mutant colonies are white, but the wild type *ADE3* on the plasmid complements *ade3* and turns the cells red. A mutant (*d-*) that requires the plasmid for viability will form a uniformly red and 5FOA-sensitive colony.

**Table S2.**
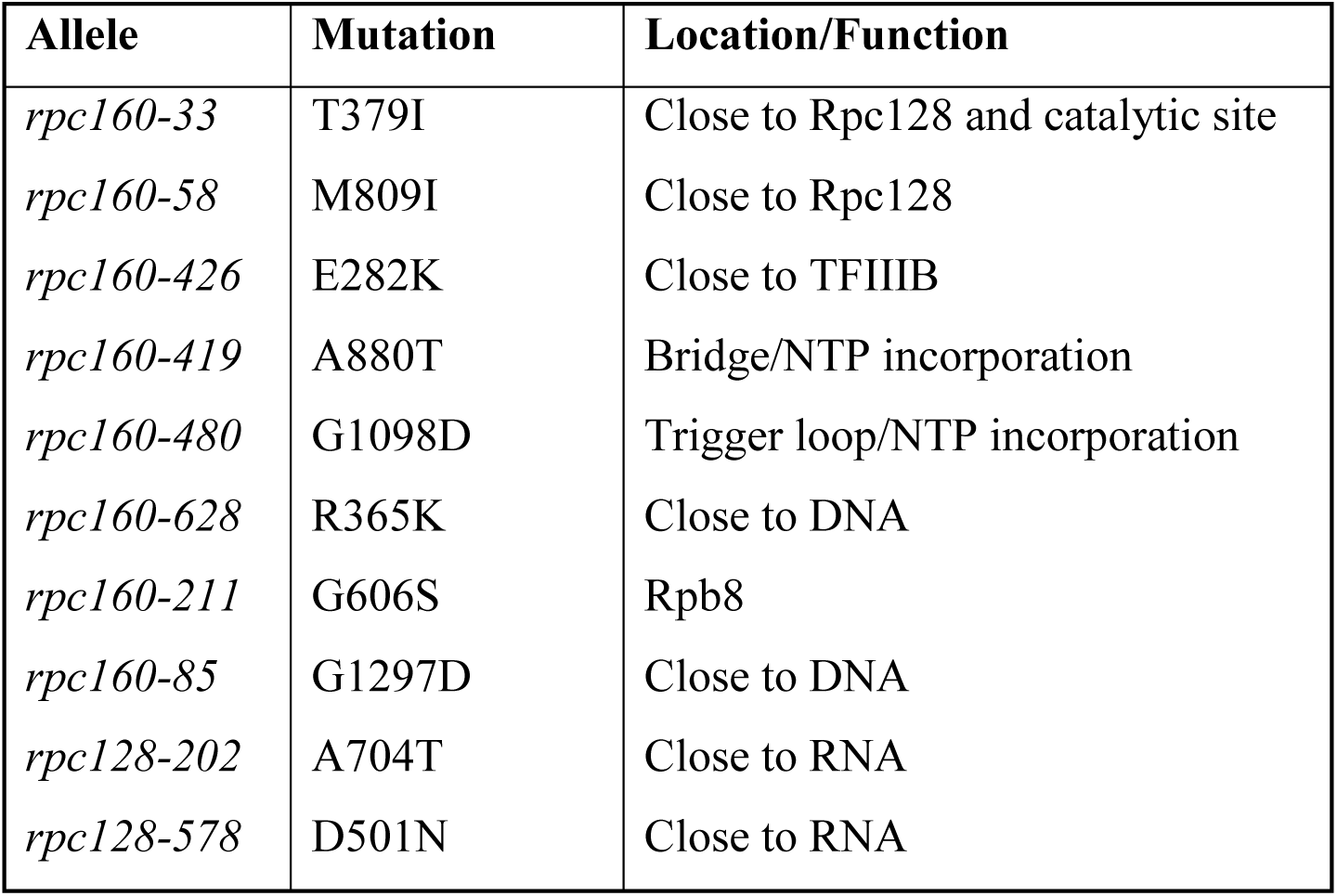
Position of the mutated residues in Pol III structure.

**Table S3.** Growth phenotype of Pol III disease mutations in yeast. *CEN LEU2* plasmids containing the indicated mutant alleles (e.g. *rpc160* mutants) were transformed into a corresponding null strain in wild type *SIZ1* or *siz1Δ* background (e.g. *rpc160A* and *rpc160A siz1Δ* strains) covering by a wild type *URA3* plasmid (e.g. a *URA3 RPC160* plasmid). Transformants were then patched on a SC-Leu plate, replica plated to 5FOA plates, and incubated at 30°C or 37°C. Growth was scored after two days of incubation. The human gene names were shown in parenthesis underneath the yeast gene names.

| Gene | Mutation in human protein | Mutation in yeast protein | 5FOA 30°C |  | 5FOA 37°C |  |
| --- | --- | --- | --- | --- | --- | --- |
|  |  |  | *SIZ1*+ | *siz1Δ* | *SIZ1*+ | *siz1Δ* |
| *RPC160*<br>(*POLR3A*) | D372N | D384N | + | + | + | + |
|  | A387G | A399G | + | + | + | + |
|  | T553I | S565I | + | + | + | + |
|  | Q599K | Q608K | -/+ | +/- | -/+ | +/- |
|  | E644K | Q658K | + | + | + | + |
|  | G672E | G686E | + | + | + | + |
|  | C724Y | C738Y | + | + | + | + |
|  | N775I | N789I | + | + | + | + |
|  | G784S | G798S | + | + | + | + |
|  | M852V | I866V | + | + | + | + |
|  | R1005C | R1061C | + | + | + | + |
|  | R1005H | R1061H | + | + | + | + |
|  | G1240S | G1308S | + | + | + | + |
|  | E1261K | E1329K | - | - | - | - |
|  | A1331T | A1399T | + | + | + | + |
|  | D372N, N775I | D384N, N789I | - | +/- | - | -/+ |
|  | Q599K, G1240S | Q608K, G1308S | -/+ | +/- | - | - |
| *RPC128*<br>(*POLR3B*) | T503K | T521K | + | + | + | + |
|  | V523E | I541E | + | + | + | + |
|  | D895N | D910N | + | + | + | + |
|  | L1012P | L1027P | -/+ | + | - | +/- |
|  | L1117V | L1132V | + | + | + | + |
|  | V523E, D895N | I541E, D910N | - | - | - | - |
|  | V523E, L1012P | I541E, L1027P | - | - | - | - |
|  | V523E, L1117V | I541E, L1132V | - | - | - | - |
| *BRF1*<br>(*BRF1*) | R223W | R218W | - | - | - | - |
|  | S226L | A221L | + | + | + | + |
|  | T259M | T254M | - | +/- | - | -/+ |
|  | P292H | P288H | +/- | + | -/+ | + |

**Figure S3.**
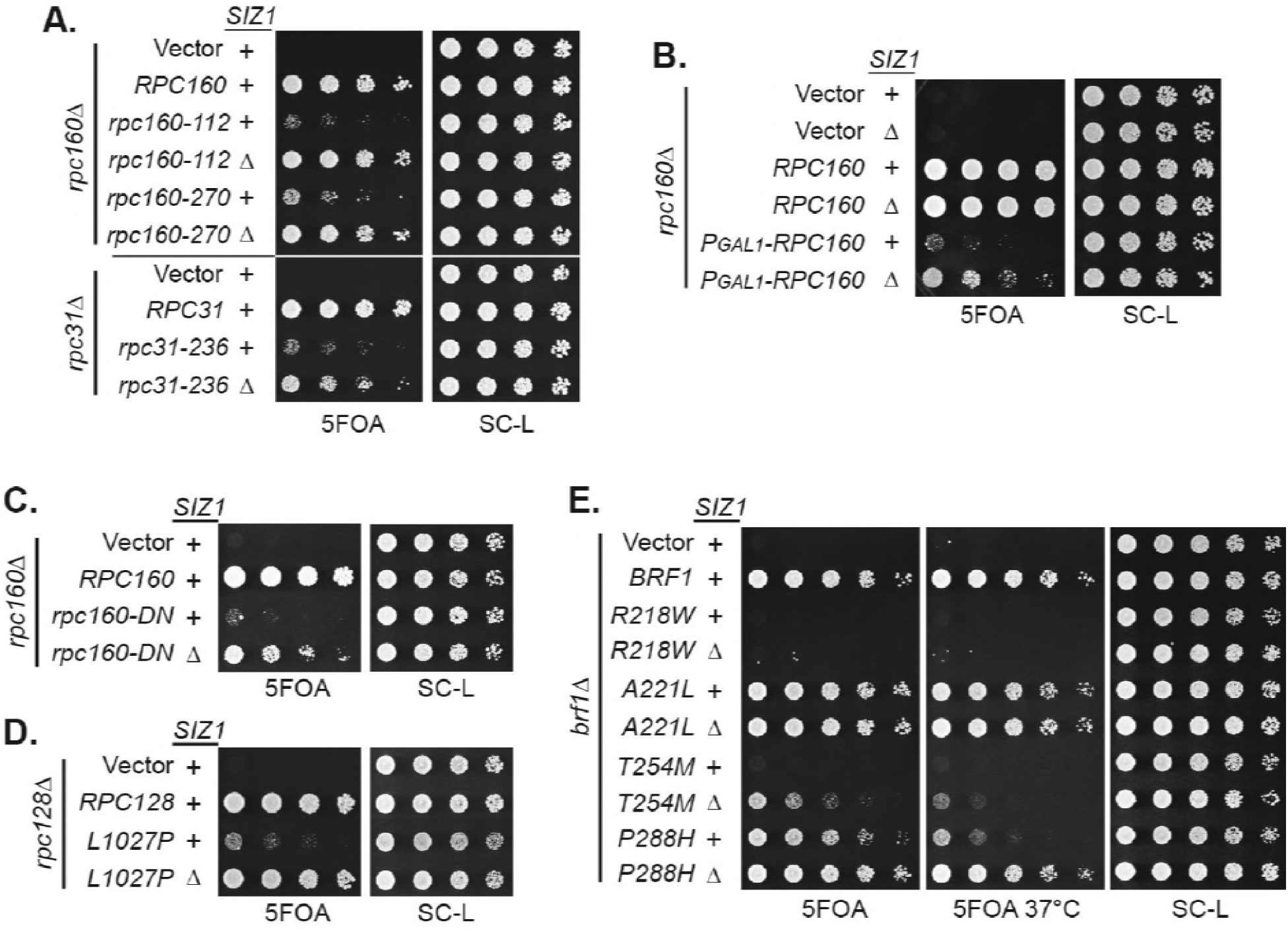
*siz1Δ* rescued a wide spectrum of Pol III mutations. **(A)** Plasmid shuffle experiments to test the growth phenotype of known *rpc160* and *rpc31* mutants in wild type *SIZ1* or *siz1A* background. The *RPC160* or *RPC31* alleles were on *LEU2* vectors. The parental *rpc160Δ* and *rpc31Δ* strains contain a *URA3* plasmid carrying wild type *RPC160* or *RPC31*, respectively. **(B)** Wild type *RPC160* was placed under control of the *GAL1* promoter on *a LEU2* vector then transformed into an *rpc160* strain carrying a *URA3 RPC160* plasmid, which was lost in the presence of 5FOA. The 5FOA plate contained glucose as the only carbon source, which strongly repressed *RPC160* expression, making the cells grow extremely slowly. *siz1Δ* partially alleviated this growth defect. **(C-E)** Similar plasmid shuffle experiments showing the rescue effect *of siz1Δ* on disease causing mutations. *rpc160-DN:* D384N, N789I.

**Figure S4.**
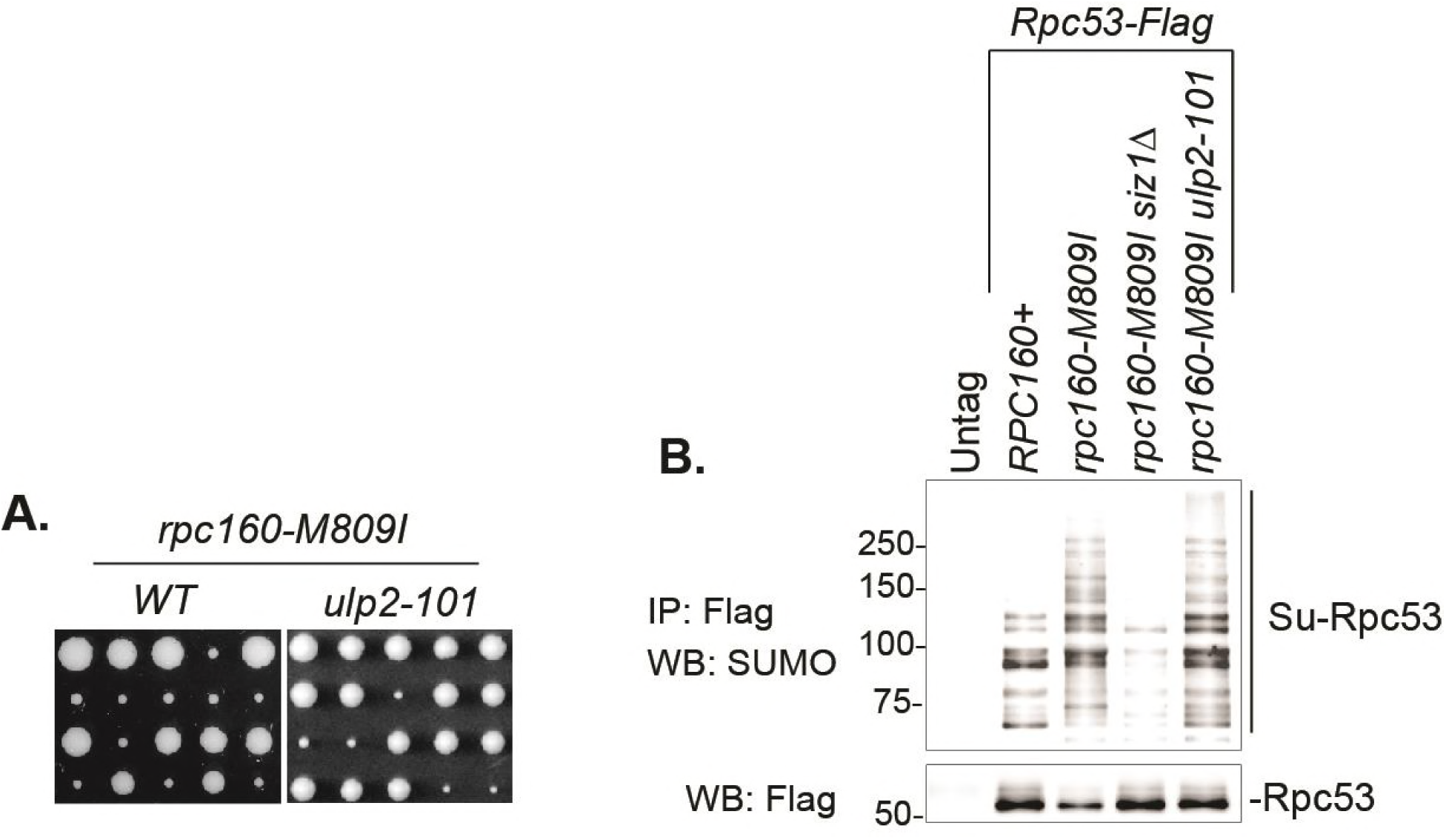
*ulp2-101* rescued *rpc160-M809I* without abolishing Rpc53 sumoylation. **(A)** An *rpc160-M809I* strain was crossed with a wild type (left) or an *ulp2-101* mutant strain (right), followed by tetrad analysis. The growth of five representative tetrads were shown. The cross between *rpc160-M809I* and wild type *ULP2* always yielded two large colonies and two small colonies, indicating *rpc160-M809I* caused severe growth defect. The cross between *rpc160-M809I* an*d ulp2-101* in most cases yielded three large colonies and one small colony, indicating *rpc160-M809I* was rescued by *ulp2-101.* **(B)** Flag-tagged Rpc53 proteins from the indicated strains were purified using anti-Flag beads, followed by SDS-PAGE and immunoblotting using an anti-SUMO antibody (top) or an anti-Flag antibody (bottom). *siz1Δ* abolished Rpc53 sumoylation, while *ulp2-101* did not.

**Table S4.**
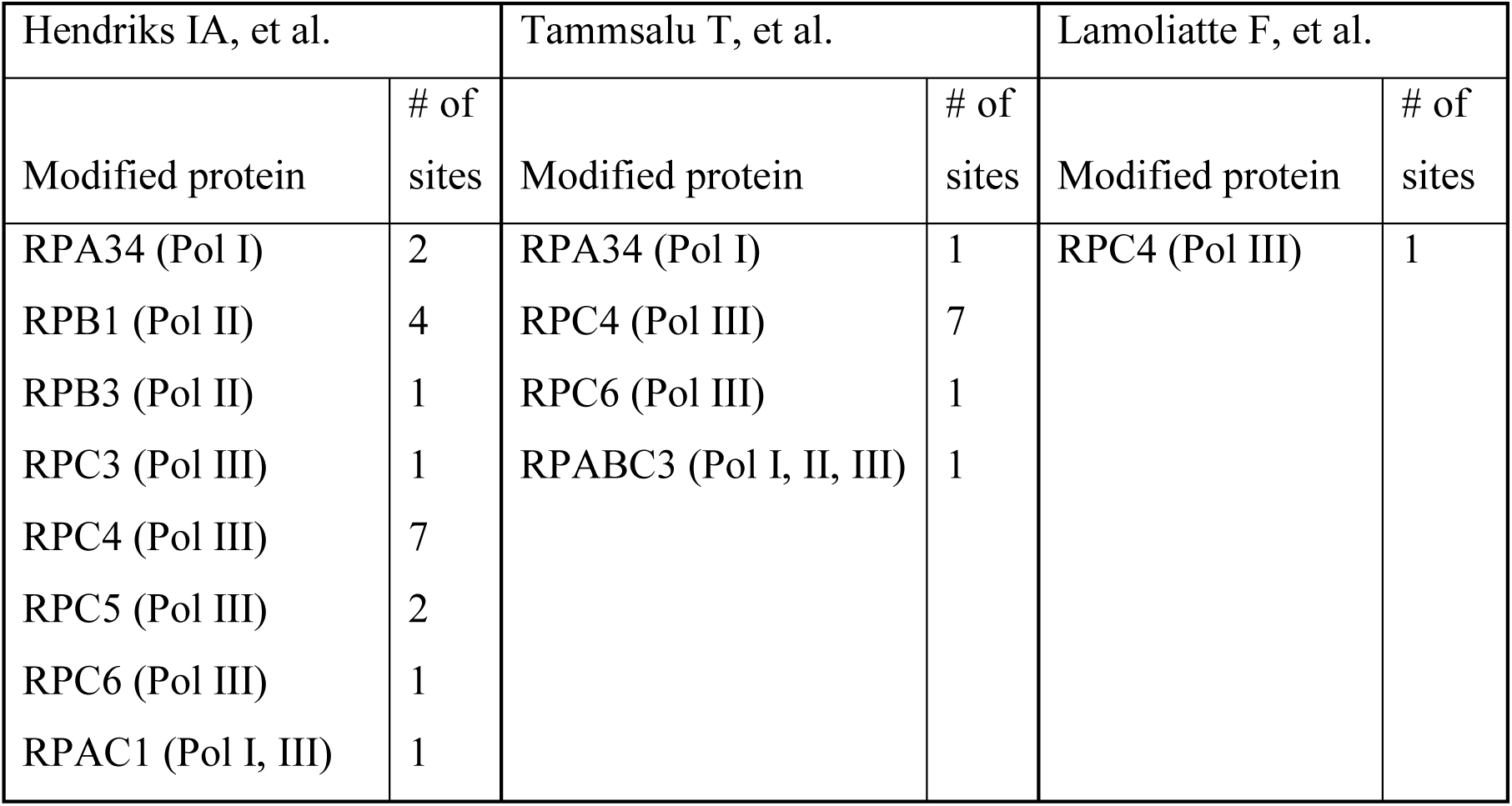
Summary of sumoylated polymerase subunits

| Hendriks IA, et al. |  | Tammsalu T, et al. |  | Lamoliatte F, et al. |  |
| --- | --- | --- | --- | --- | --- |
| Modified protein | # of sites | Modified protein | # of sites | Modified protein | # of sites |
| RPA34 (Pol I) | 2 | RPA34 (Pol I) | 1 | RPC4 (Pol III) | 1 |
| RPB1 (Pol II) | 4 | RPC4 (Pol III) | 7 |  |  |
| RPB3 (Pol II) | 1 | RPC6 (Pol III) | 1 |  |  |
| RPC3 (Pol III) | 1 | RPABC3 (Pol I, II, III) | 1 |  |  |
| RPC4 (Pol III) | 7 |  |  |  |  |
| RPC5 (Pol III) | 2 |  |  |  |  |
| RPC6 (Pol III) | 1 |  |  |  |  |
| RPAC1 (Pol I, III) | 1 |  |  |  |  |

**Table S5.**
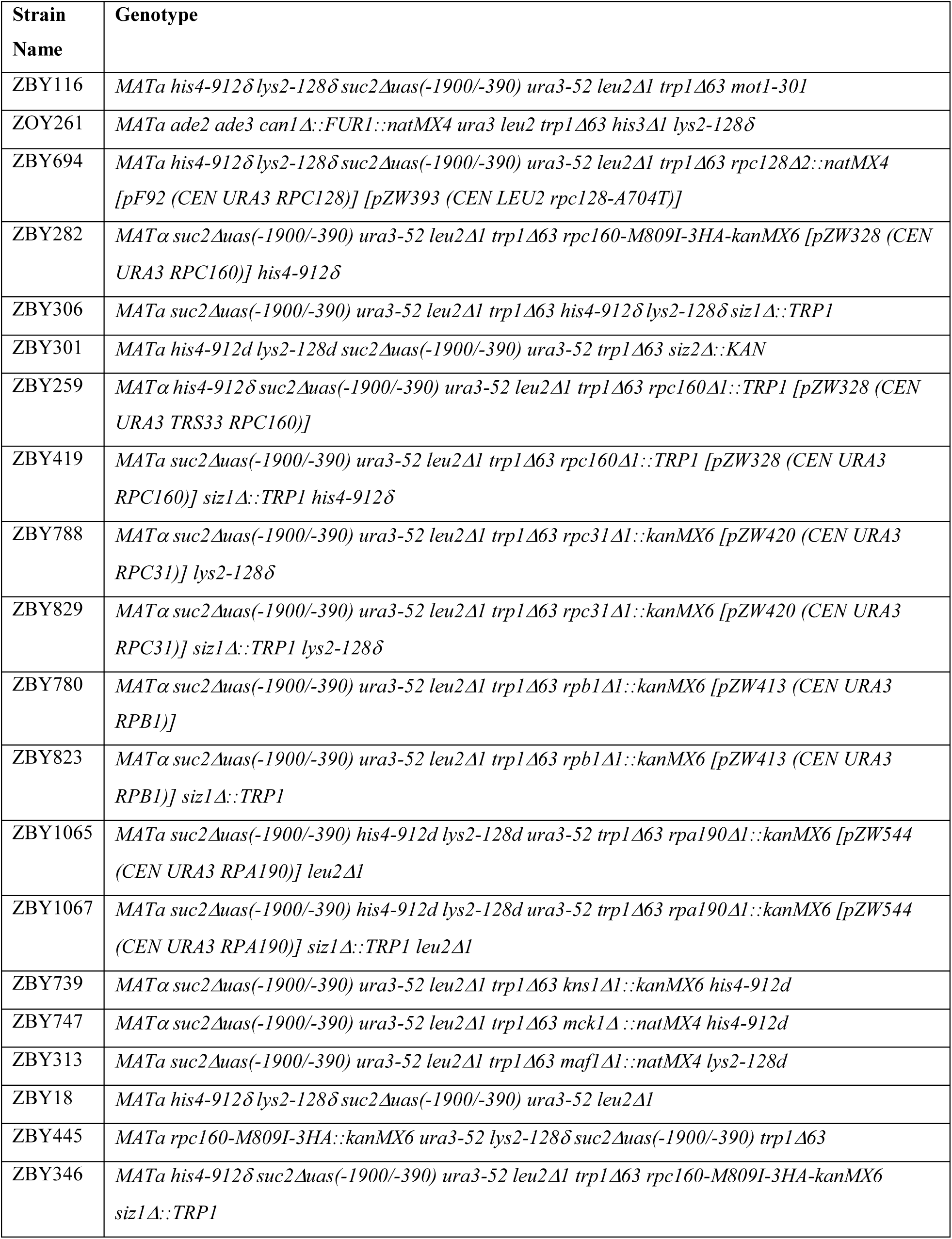
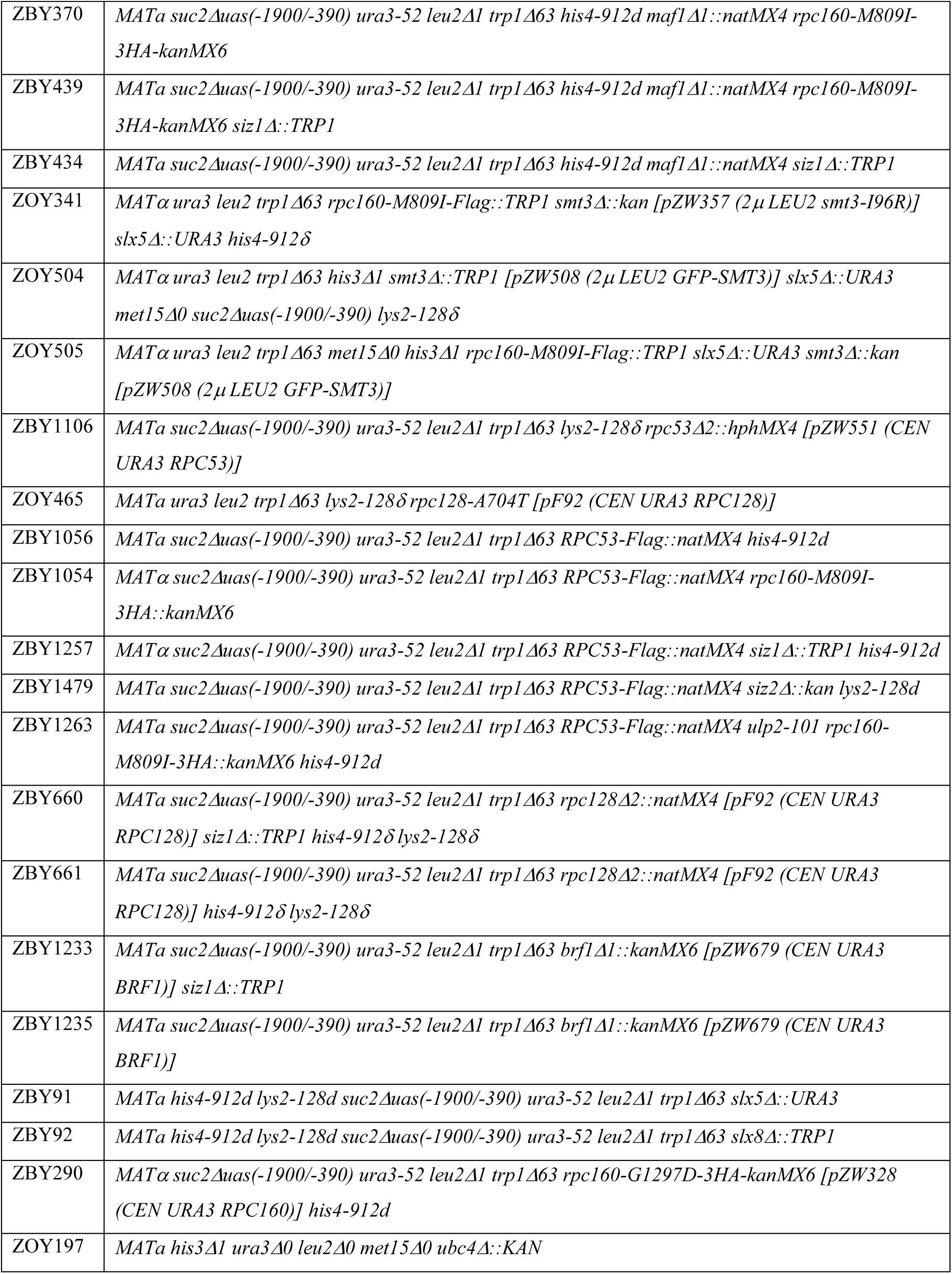
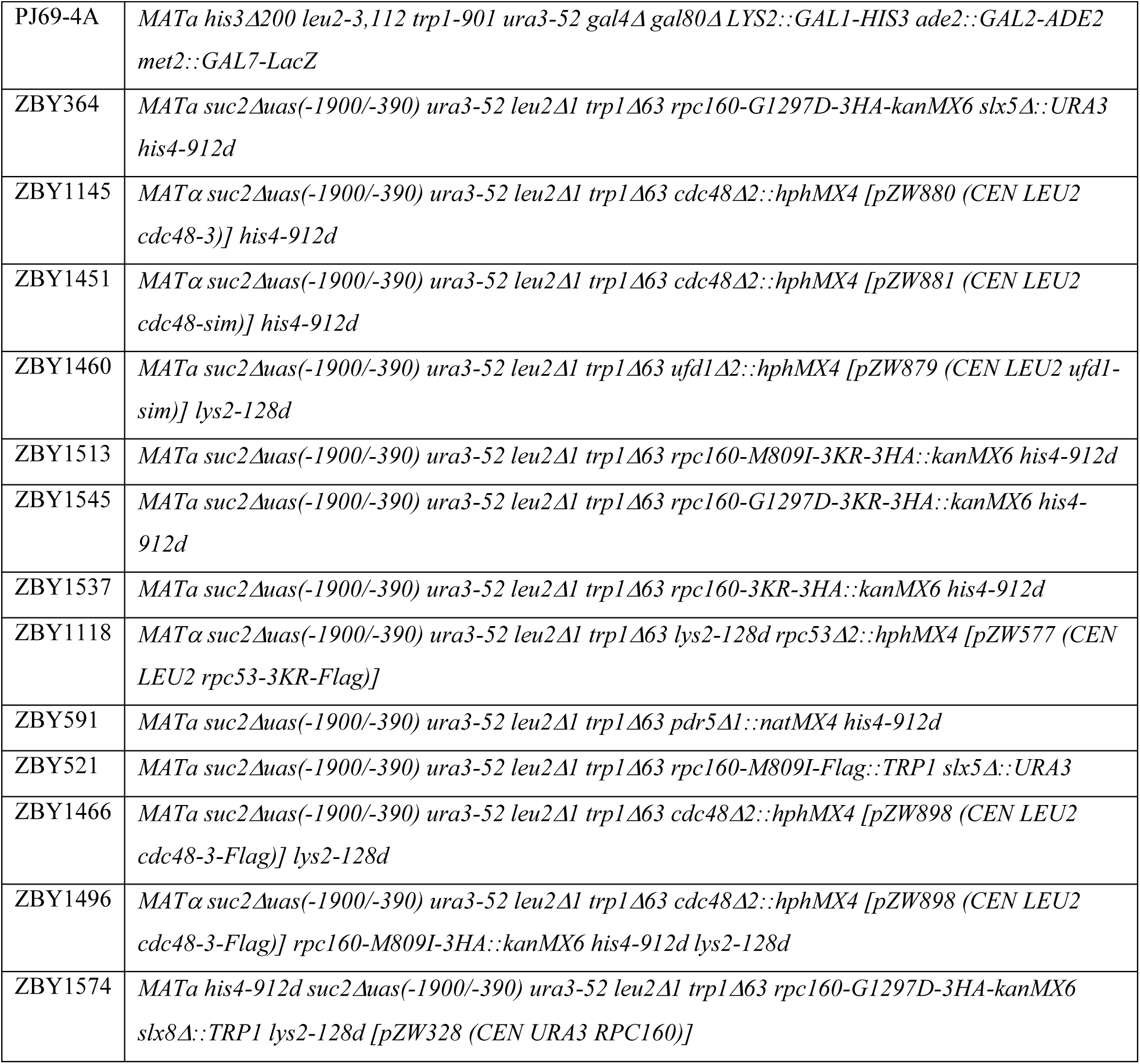
Yeast strains used in this study.

**Table S6.** Plasmids used in this study.

| Plasmid name | Markers |
| --- | --- |
| pRS415 | *Amp CEN LEU2* |
| pRS425 | *Amp 2μ LEU2* |
| pF5 | *Amp CEN LEU2 SMT3* |
| pZW81 | *Amp CEN LEU2 MOT1* |
| pZW311 | *Amp CEN LEU2 SMT3-Q56K* |
| pZW321 | *Amp CEN URA3 ADE3 SMT3-Q56K* |
| pAK21-1 | *Amp CEN TRP1 ade3-pink* |
| pZW548 | *Amp CEN LEU2 RPA190* |
| pZW890 | *Amp CEN LEU2 rpa190-G1194D* |
| pZW411 | *Amp CEN LEU2 RPB1* |
| pZW8910 | *Amp CEN LEU2 rpb1-G1073D* |
| pZW326 | *Amp CEN LEU2 RPC160* |
| pZW371 | *Amp CEN LEU2 rpc160-G1098D* |
| pZW400 | *Amp CEN LEU2 rpc160-112* |
| pZW401 | *Amp CEN LEU2 rpc160-270* |
| pZW873 | *Amp CEN LEU2 P<sub>GAL1</sub>-RPC160-Flag* |
| pZW419 | *Amp CEN LEU2 RPC31* |
| pZW423 | *Amp CEN LEU2 rpc31-236* |
| pZW725 | *Amp CEN LEU2 RPC53* |
| pZW564 | *Amp CEN LEU2 RPC53-Flag* |
| pZW565 | *Amp CEN LEU2 rpc53-K51R-Flag* |
| pZW566 | *Amp CEN LEU2 rpc53-K115R-Flag* |
| pZW567 | *Amp CEN LEU2 rpc53-K216R-Flag* |
| pZW569 | *Amp CEN LEU2 rpc53-K236R-Flag* |
| pZW570 | *Amp CEN LEU2 rpc53-K115,236R-Flag* |
| pZW575 | *Amp CEN LEU2 rpc53-K51,115R-Flag* |
| pZW576 | *Amp CEN LEU2 rpc53-K51,236R-Flag* |
| pZW577 | *Amp CEN LEU2 rpc53-K51,115,236R-Flag* |
| pZW568 | *Amp CEN LEU2 rpc53-K322,325R-Flag* |
| pZW571 | *Amp CEN LEU2 rpc53Δ2-275-Flag* |
| pZW384 | *Amp CEN LEU2 RPC128* |
| pZW596 | *Amp CEN LEU2 smt3ΔGG-rpc53-K51,115,236R-Flag* |
| pZW630 | *Amp CEN LEU2 rpc160-S565I* |
| pZW579 | *Amp CEN LEU2 rpc160-D384N, N789I* |
| pZW640 | *Amp CEN LEU2 rpc160-E1329K* |
| pZW387 | *Amp CEN LEU2 rpc128-I541E* |
| pZW624 | *Amp CEN LEU2 rpc128-L1027P* |
| pZW678 | *Amp CEN LEU2 BRF1* |
| pZW680 | *Amp CEN LEU2 brf1-R218W* |
| pZW681 | *Amp CEN LEU2 brf1-A221L* |
| pZW682 | *Amp CEN LEU2 brf1-T254M* |
| pZW683 | *Amp CEN LEU2 brf1-P288H* |
| pZW144 | *Amp 2μ LEU2 SMT3* |
| pGADT7 | *Amp 2μ LEU2GAL4-AD* |
| pGBKT7 | *Kan 2μ TRP1 GAL4-BD* |
| pCS6514 | *Amp 2μ LEU2 GAL4-AD-SLX5* |
| pZW584 | *Kan 2μ TRP1 GAL4-BD-RPC53* |
| pZW591 | *Kan 2μ TRP1 GAL4-BD-rpc53-3KR* |
| pZW592 | *Kan 2μ TRP1 GAL4-BD-rpc53ΔN (Δ2-275)* |
| pZW331 | *Amp CEN URA3 RPC160-HA* |
| pZW332 | *Amp CEN URA3 rpc160-M809I-HA* |
| pZW968 | *Amp CEN URA3 rpc160-M809I-3KR-HA* |
| pZW938 | *Amp CEN LEU2 SIZ1-Myc* |
| pZW940 | *Amp CEN LEU2 PCUP1-siz1ΔSAP-Myc (Δ34-68)* |
| pZW941 | *Amp CEN LEU2 PCUP1-siz1-SAP\*-Myc* |
| pGP776 | *Amp CEN LEU2 SIZ1* |
| pZW903 | *Amp CEN LEU2 SIZ1-Flag* |
| pZW904 | *Amp CEN LEU2 PCUP1-siz1ΔSAP-Flag* |
| pZW942 | *Amp CEN LEU2 PCUP1-siz1-SAP\*-Flag* |
| pZW986 | *Amp 2μ LEU2 HA-SLX8* |
| pZW993 | *Amp 2μ LEU2 P<sub>CUP1</sub>-HA-slx8ΔN (Δ2-163)* |

**Table S7.**
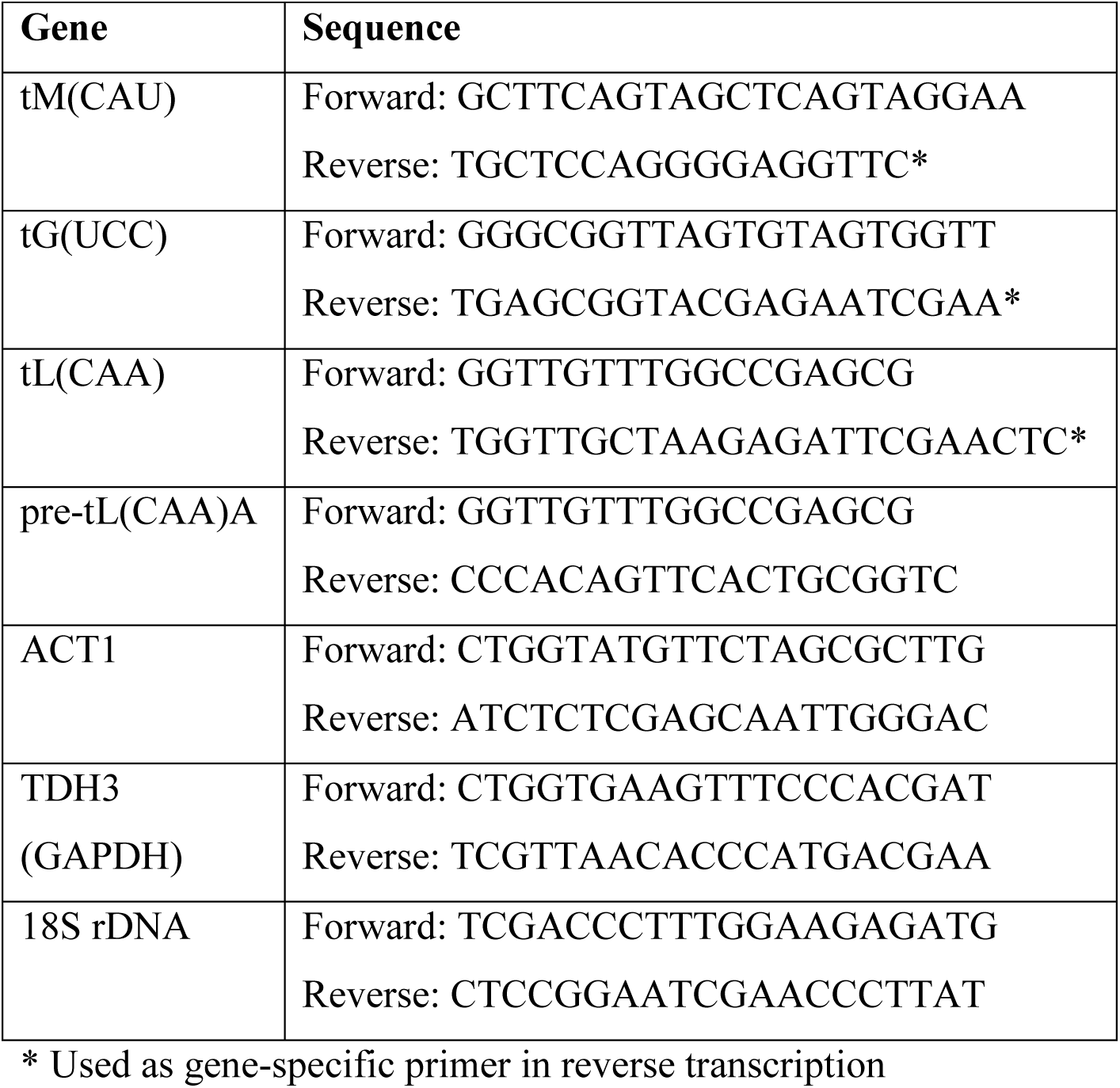
Primers used in this study. The same primers were used in RNA level measurement and in chromatin IP experiments.

